# Insulin-Independent Regulation of Type 1 Diabetes via Brown Adipocyte-Secreted Proteins and the Novel Glucagon Regulator Nidogen-2

**DOI:** 10.1101/2024.08.30.610490

**Authors:** Jeongmin Lee, Alessandro Ustione, Emily M Wilkerson, Rekha Balakrishnan, Debbie C. Thurmond, Dennis Goldfarb, David W. Piston

**Affiliations:** Department of Cell Biology & Physiology, Washington University School of Medicine, St. Louis, Missouri, United States of America; Department of Molecular & Cellular Endocrinology, City of Hope, Duarte, California, United States of America

**Author notes:** Corresponding author. (DWP).

## Abstract

Current treatments for type 1 diabetes (T1D) focus on insulin replacement. We demonstrate the therapeutic potential of a secreted protein fraction from embryonic brown adipose tissue (BAT), independent of insulin. The large molecular weight secreted fraction mediates insulin receptor-dependent recovery of euglycemia in a T1D animal model, nonobese diabetic (NOD) mice, by suppressing glucagon secretion. This fraction also promotes white adipocyte differentiation and browning, maintains healthy BAT, and enhances glucose uptake in adipose tissue, skeletal muscle, and liver. From this fraction, we identify nidogen-2 as a critical BAT-secreted protein that reverses hyperglycemia in NOD mice, inhibits glucagon secretion from pancreatic α-cells, and mimics other actions of the entire secreted fraction. These findings confirm that BAT transplants affect physiology and demonstrate that BAT-secreted peptides represent a novel therapeutic approach to diabetes management. Furthermore, our research reveals a novel signaling role for nidogen-2, beyond its traditional classification as an extracellular matrix protein.

**HIGHLIGHTS:**

- The large molecular weight brown adipocyte-secreted protein fraction suppresses glucagon secretion and normalizes glycemia in mouse models of type 1 diabetes (T1D), independent of insulin, offering a novel therapeutic strategy for disease management.
- Nidogen-2, a critical component of this fraction, is identified as an inhibitor of glucagon secretion in pancreatic α-cells by regulating intracellular messenger activities.
- The large-secreted protein fraction prevents T1D-related whitening of brown adipose tissue, promotes adipocyte differentiation, and enhances browning of inguinal white adipose tissue.
- This fraction enhances glucose uptake in adipose tissue, skeletal muscle, and liver through an insulin receptor-dependent pathway.

## INTRODUCTION

Type 1 diabetes (T1D) is defined by autoimmune-mediated destruction of the insulin-producing pancreatic β-cells and exacerbated by the aberrant hypersecretion of glucagon by α-cells ^1^. Current therapeutic strategies for T1D focus on insulin replacement through subcutaneous administration of the hormone or islet/pancreas transplantation ^2^. These methods are effective but imperfect since exogenous insulin administration cannot replicate physiologic insulin release, and transplantation requires immunosuppression ^2,3^. Consequently, there is a continuing need to discover and validate additional tools for T1D therapy ^3,4^.

Brown adipose tissue (BAT) has gained attention in metabolic research due to its endocrine functions and role in metabolic regulation ^5,6^. Beyond its well-documented capacity for thermogenesis, BAT acts as a secretory organ, modulating metabolic processes through autocrine, paracrine, and endocrine pathways ^7^. BAT-secreted extracellular vesicles (EVs), proteins, and non-protein factors influence a wide array of metabolic functions, including BAT differentiation, metabolic health, and glucose and lipid homeostasis ^7,8^. Given its profound impact on metabolism, BAT transplantation has been explored as a novel therapeutic intervention for ameliorating glucose dysregulation in metabolic disorders like obesity and diabetes ^9–12^.

Subcutaneous BAT transplantation improved body weight control, glucose metabolism, and cardiac health in type 2 diabetes mouse models ^9^. The transplantation of embryonic BAT (eBAT) into T1D mouse model has also shown insulin-independent recovery of euglycemia, accompanied by elevation of IGF-1 and suppression of glucagon secretion ^13,14^. Here, we demonstrate that secreted proteins from BAT mediate the positive effects of transplantation by showing that injection of fractionated BAT-secreted proteins improves glucose homeostasis. We identify nidogen-2 as a key component of the active fraction and show that this protein alone mimics many of the effects of tissue transplants. These data point to fresh therapeutic strategies for T1D by leveraging BAT-secreted proteins, including nidogen-2, to improve whole-body glucose metabolism.

## RESULTS

### Characterization of Large Brown Adipocyte-Secreted Proteins (CB-100)

Previous studies have demonstrated a decrease in plasma glucagon levels following the transplantation of eBAT into T1D mouse model (both STZ-induced and nonobese diabetic, NOD mice) ^13–15^. Further, conditioned buffer from the brown adipocyte cell line (nbat9) inhibited glucagon secretion from pancreatic islets ^16^. We analyzed various properties of the brown adipocyte-conditioned buffer to identify the factor responsible for the glucagon secretion inhibition.

Suppression of glucagon secretion in non-diabetic murine islets is seen in low glucose conditions with conditioned buffer from immortalized brown adipocytes (imBAT) with size fractions exceeding 100 kDa, as well as those within the 1 kDa to 3 kDa range (Figure 1A). Since the size fraction larger than100kDa exhibits the strongest suppressive effect, we chose this size fraction (conditioned buffer exceeding molecular size of 100kDa; termed CB-100) for further investigation. CB-100 was collected from incubating adult BAT (aBAT), eBAT, and cultured imBAT cells, and tested for glucagon inhibition activity. Glucagon secretion-suppressing activity is observed with eBAT and imBAT but not aBAT (Figure 1B). These results suggest that the glucagon secretion inhibiting CB-100 is more highly secreted in the embryonic stage of BAT compared to mature BAT, which is consistent with previously reported suppression of hyperglucagonemia in T1D with eBAT transplantation but not aBAT ^13^. Given the similarity of glucagon secretion inhibition by CB-100 from imBAT to that from eBAT, imBAT was utilized to collect CB-100 on a large scale for subsequent experiments. CB-100 suppresses glucagon secretion *ex vivo* from both mouse and human islets in low glucose settings, regardless of their diabetic status (Figures 1C and 1D). However, CB-100 does not affect insulin secretion in either mouse or human pancreatic islets (Figure S1A).

**Figure 1.**
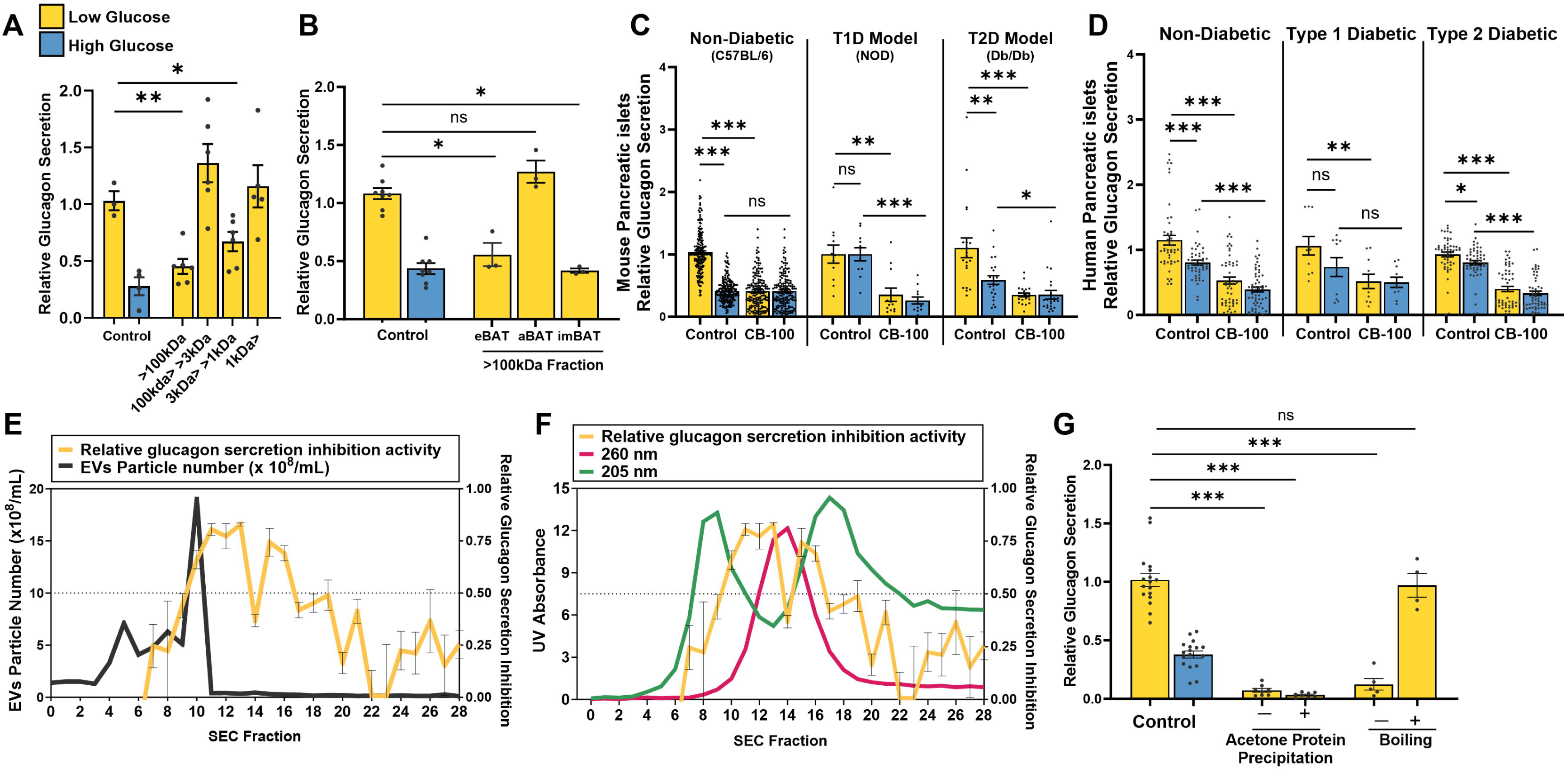
Characterization of Large Brown Adipocyte-Secreted Proteins (CB-100) (**A**) Relative *ex vivo* glucagon secretion measured in non-diabetic mouse islets (n=3) with different molecular size fractions from an immortalized brown adipocyte cell line (imBAT) conditioned buffer (larger than 100kDa, between 100kDa and 3kDa, between 3kDa and 1kDa, smaller than 1kDa; 20 μg /mL) at low (yellow, 1mM) glucose condition. (**B**) Relative glucagon secretion measured in non-diabetic mouse islets (n=4) with >100kDa fraction from embryonic (eBAT, E14-16, n=6), adult (aBAT, n=4) adipose tissue, and imBAT (20 μg/mL) at low (yellow, 1mM) glucose condition. **(C&D)** Relative glucagon secretion measured in low (yellow, 1mM) and high (teal, 11mM) glucose conditions in both non-diabetic and diabetic (**C**) mouse (non-diabetic n=30, T1D Model (NOD) n=5, T2D (Db/Db) model n=3) and **(D**) human (non-diabetic n=16, T1D n=1, T2D n=8, see Table S1 for more detail) pancreatic islets. (**E**) Relative glucagon secretion inhibition activity measured at low (1mM) glucose in non-diabetic mouse islets (n=5) (yellow) plotted with extracellular vesicles (EVs) particles number (black) of size exclusion column (CL-2B) fractions with CB-100. (**F**) Relative glucagon secretion inhibition activity measured at low (1mM) glucose in non-diabetic mouse islets (n=5) (yellow) plotted with 205nm (green) and 260nm (red) absorbance of size exclusion column (CL-2B) fractions with CB-100. (**G**) Relative glucagon secretion measured in non-diabetic mouse islets (n=3) with CB-100 with or without acetone protein precipitation or boiling at low (yellow, 1mM) glucose conditions. Data are presented as Mean ± SEM. Statistical significance was determined using an unpaired t-test. Groups compared for statistical analysis are indicated by the line. *p < 0.05, **p < 0.01, ***p < 0.001, ns (p > 0.05).

Since a hallmark of brown adipocytes is thermogenesis, we tested if the glucagon secretion-inhibiting properties of CB-100 are related to thermogenic activity of imBAT. Norepinephrine (NE) stimulation increases mRNA expression of the thermogenic activity marker genes (uncoupling protein 1 (*Ucp1*) and cell death Inducing DFFA like effector A (*Cidea*)) in imBAT (Figure S1B), but the glucagon secretion inhibiting activity shows no difference between CB-100 from NE stimulated and unstimulated imBAT (Figure S1C).

Adipose tissue-secreted EVs have also been recognized as important players in intercellular and organ crosstalk ^17^. We investigated whether the glucagon secretion-inhibiting activity of CB-100 originated from imBAT-secreted EVs. The number of EVs particles does not strongly correlate with the glucagon secretion inhibition activity (Figure 1E). Instead, peptide bond absorbance (205 nm and 260 nm) shows a better correlation (Figure 1F), suggesting that the CB-100-mediated suppression of glucagon secretion is mediated by large proteins (>100 kDa) rather than through EVs. To confirm the protein-like property of the glucagon secretion-inhibiting factor, proteins were isolated from CB-100 by acetone precipitation and heated by boiling. The glucagon secretion-suppressing activity was preserved through acetone protein precipitation but abolished by boiling (Figure 1G), indicating that the glucagon secretion-inhibiting factors within CB-100 exhibit biochemical characteristics of proteins.

### CB-100 Enhances Glucose Management in a Type 1 Diabetes Model

The *in vivo* impact of CB-100 was investigated using NOD mice as a T1D model. Upon reaching the early hyperglycemic stage (between 190mg/dL and 230mg/dL), NOD mice received daily subcutaneous injections of CB-100 (1.5 mg/kg body weight) or sham (KRBH buffer) for seven consecutive days. After one week, the injections were stopped, and animals’ blood glucose levels were monitored weekly for 2∼8 weeks. Subsequently, metabolic activity, glucose and insulin tolerance were measured, and post-mortem organ analyses were conducted (Figure 2A). Regardless of sex and age, 30 out of 33 CB-100-treated mice (19 female and 11 male) exhibit a significant and sustained reduction in blood glucose levels for four to nine weeks, while sham-treated mice develop severe hyperglycemia (>600mg/dL) within two weeks (Figures 2B, S2B and 2E). Administering the treatment during the early hyperglycemia resulted in a higher success rate compared to the later stage (blood glucose levels >230mg/dL, with a success rate of 21.1%) (Figures S2A, S2C, S2D and S2F).

**Figure 2.**
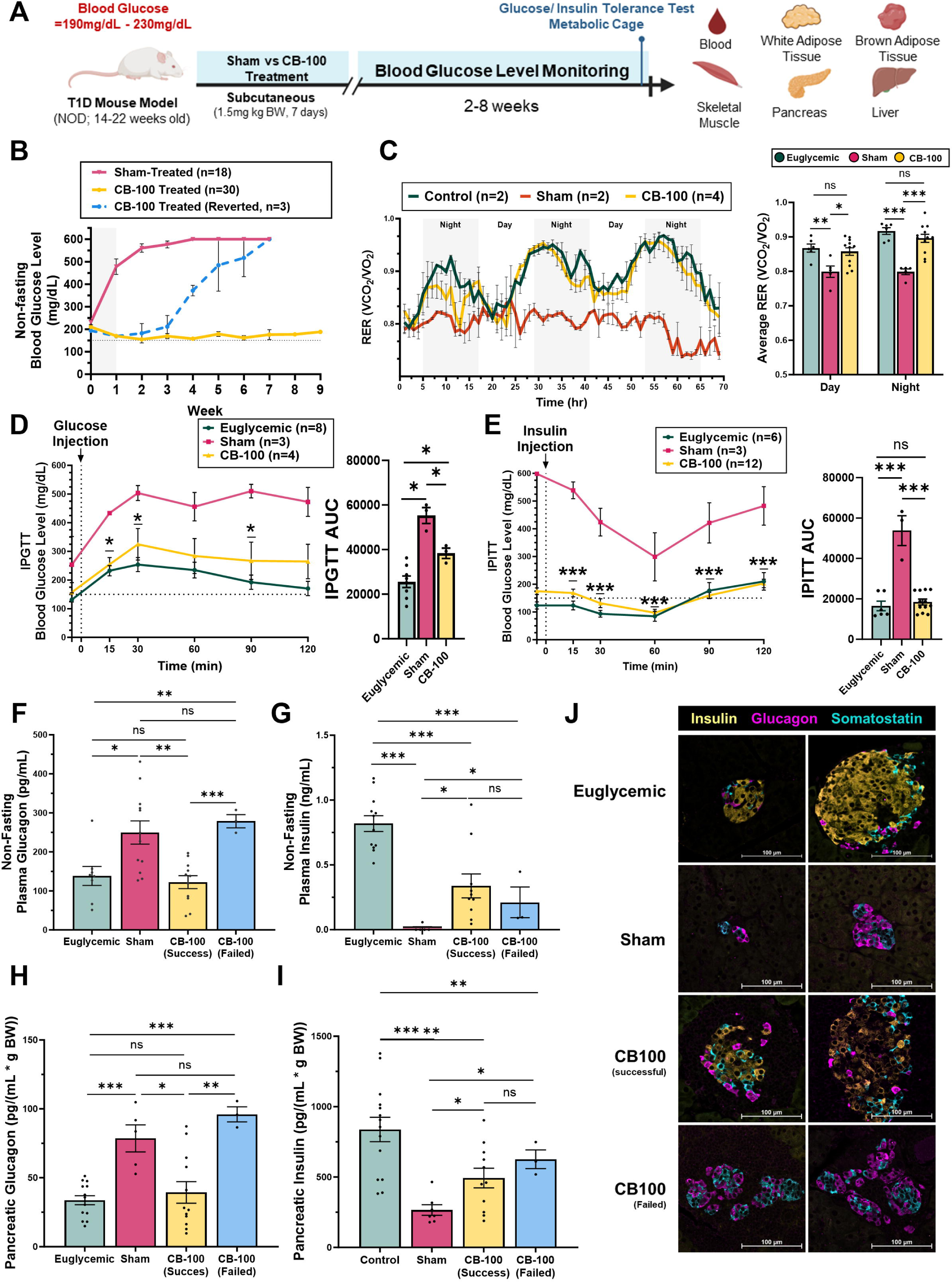
CB-100 Enhances Glucose Management in a Type 1 Diabetes Model by Reversing Hyperglucagonemia. **(A)** Schematic of the experimental design for CB-100 injection in nonobese diabetic (NOD) mice (14-22weeks old); Once NOD mice reached the early stage of hyperglycemia (blood glucose level between 190mg/dL and 230mg/dL), 1.5mg/kg body weight (BW) CB-100 subcutaneous injection was done for 7 days. Afterward, weekly blood glucose monitoring was performed using an intraperitoneal glucose tolerance test (IPGTT), an intraperitoneal insulin tolerance test (IPITT), and metabolic cages studies. After 2 to 8 weeks of monitoring, mice were euthanized, and postmortem tissues (blood, adipose tissue, skeletal muscle, pancreas, and liver) were collected for further analysis. **(B)** Weekly non-fasting blood glucose levels measured following 7 days of injection of CB-100 (subcutaneous, 1.5 mg/kg BW, successful; yellow, n = 30 (19 females, 11 males), reverted; blue, n = 3 (2 females, 1 male)) and sham (KRBH, red; n = 18 (male and female)) in NOD mice (14-22 weeks old). Normal non-fasting blood glucose level for NOD mice was indicated with a dotted line (150mg/dL). Grey highlighted days are the days of the CB-100 injections. **(C)** Respiratory Exchange Rate (RER) measured in CB-100-treated (yellow, n=4), sham (red, n = 2), and euglycemic control (green, n=2) NOD mice (female). Measurements were taken every 15 minutes over 3 days, with four measurements averaged for each hourly data point. See supplementary Fig S2 G-I for more detailed metabolic profiling. **(D)** IPGTT blood glucose levels measured at 15, 30, 60, 90, and 120 minutes post 2g/kg BW glucose administration to euglycemic control (green, n=8), CB-100-treated (yellow, n=4), and sham-treated (red, n=3) female NOD mice and calculated Area under the curve (AUC). NOD mouse normal non-fasting blood glucose level is marked with a dotted line (∼150mg/dL). **(E)** IPITT blood glucose levels measured at 15, 30, 60, 90, and 120 minutes post 0.75 IU/kg BW insulin administration to euglycemic control (green, n=6), CB-100 treated (yellow, n=12), sham-treated (red, n=3) female NOD mice and calculated AUC. NOD mouse normal non-fasting blood glucose level is marked with a dotted line (150mg/dL). **(F)** Non-fasting blood plasma glucagon measured in euglycemic control (green, n = 8), sham-treated (red, n=12), successful CB-100 treated (yellow, n=11), and failed CB-100 treated (blue, n=3) female NOD mice. **(G)** Non-fasting blood plasma insulin was measured in euglycemic control (green, n = 12), sham-treated (red, n=5), successful CB-100 treated (yellow, n=10), and failed CB-100 treated (blue, n=3) female NOD mice. **(H)** Pancreatic glucagon measured in euglycemic control (green, n = 14), sham-treated (red, n=5), successful CB-100 treated (yellow, n=12), and failed CB-100 treated (blue, n=3) female NOD mice, normalized to body weight. **(I)** Pancreatic insulin measured in euglycemic control (green, n = 14), sham-treated (red, n=7), successful CB-100 treated (yellow, n=11), and failed CB-100 treated (blue, n=3) female NOD mice, normalized to body weight. **(J)** Representative immunofluorescence images of pancreas sections for insulin (yellow), glucagon (magenta), and somatostatin (cyan) of euglycemic, sham-treated, and CB-100 treated female NOD mice. Scale bar: 100 μm. Data are presented as Mean ± SEM. Statistical significance was determined using an unpaired t-test. Groups compared for statistical analysis are indicated by the line. For IPITT and IPGTT, blood glucose level of CB-100-treated was compared with sham-treated NOD mice. *p < 0.05, **p < 0.01, ***p < 0.001, ns (p > 0.05).

Mice receiving CB-100 treatment also demonstrate improved metabolic parameters by indirect calorimetry (oxygen consumption (VCO_2_/VO_2_)) compared to the sham-treated group (Figure 2C). To evaluate the effects of CB-100 on glucose homeostasis, treated and untreated NOD mice underwent intraperitoneal glucose tolerance test (IPGTT, 2g/kg BW glucose following 6 hours fast) or intraperitoneal insulin tolerance test (IPITT, 0.75 IU/kg BW insulin following 4 hours fast). In the IPGTT, CB-100-treated NOD mice show a lower area under the curve (AUC) than sham-treated NOD mice but remain higher than the non-diabetic control mice (Figure 2D). In the IPITT, CB-100-treated mice demonstrate a lower AUC than sham-treated diabetic mice, with their AUC values similar to healthy control animals’ (Figure 2E). These findings suggest that CB-100 treatment improves glycemic control in a T1D mouse model by improving glucose tolerance and insulin sensitivity.

### CB-100-Mediated Restoration of Euglycemia Is Dependent on the Suppression of Plasma Glucagon and Is Independent of Insulin

To determine the roles of islet hormones in restoring euglycemia, we compared *in vivo* glucagon and insulin levels among non-diabetic, CB-100-treated, and sham-treated NOD mice. Compared to the sham-treatment, CB-100-treated NOD mice demonstrate a decrease in plasma glucagon levels comparable to the non-diabetic NOD mice (Figure 2F). A similar trend is observed in pancreatic glucagon contents where non-diabetic and successfully treated CB-100 NOD mice show significantly lower levels of pancreatic glucagon compared to sham-treated and unsuccessful CB-100-treated NOD mice (Figure 2H). Insulin levels do not exhibit similar patterns. In both plasma and pancreas, insulin levels are lower for CB-100 treated mice than for non-diabetic NOD mice (Figures 2G and 2I). While CB-100-treated NOD mice exhibit higher insulin levels than the sham-treated mice, these levels are both significantly lower than non-diabetic control NOD mice. There was no significant difference in insulin levels between successfully and unsuccessfully CB-100-treated mice (Figures 2G and 2I), which suggests that insulin levels are not a major determining factor for restoring euglycemia in this context.

Histological analysis reveals insulin-positive cells in pancreata from CB-100-treated NOD mice, though these cells are not as abundant as in non-diabetic controls (Figure 2J). Additionally, successfully treated CB-100 mice exhibit reduced α-cell hyperplasia, showing a lower ratio of glucagon-positive cells relative to islet size than failed and sham-treated NOD mice (Figure 2J). Unlike the plasma and pancreatic hormone measurements, pancreata from mice with failed CB-100 treatments do not show insulin-positive cells.

### CB-100 Enhances the Adipocyte Differentiation and Browning in White Adipose Tissue

Given the critical role of adipose tissue in glucose uptake ^18^, we analyzed inguinal white adipose tissue (ingWAT) in NOD mice. Since *in vivo* browning of ingWAT has been reported ^19^, we assessed the effects of CB-100 treatment on ingWAT and its potential contributions to glucose homeostasis. CB-100-treated mice show an increased body weight compared to sham-treated mice, but they do not exceed the percentage weight change of non-diabetic control mice before and after CB-100 treatment (Figure 3A). Body composition analysis reveals a higher percentage of fat mass in CB-100-treated mice compared to the sham-treated, with no significant differences in lean mass (Figure 3B). Normalizing ingWAT mass to total body weight, CB-100-treated mice display increased ingWAT mass compared to sham-treated NOD mice (Figure 3C). Hematoxylin and eosin (H&E) staining of control and CB-100 treated ingWAT show unilocular lipid droplets, while sham-treated diabetic controls show adipose tissue fibrosis (Figure 3H).

**Figure 3.**
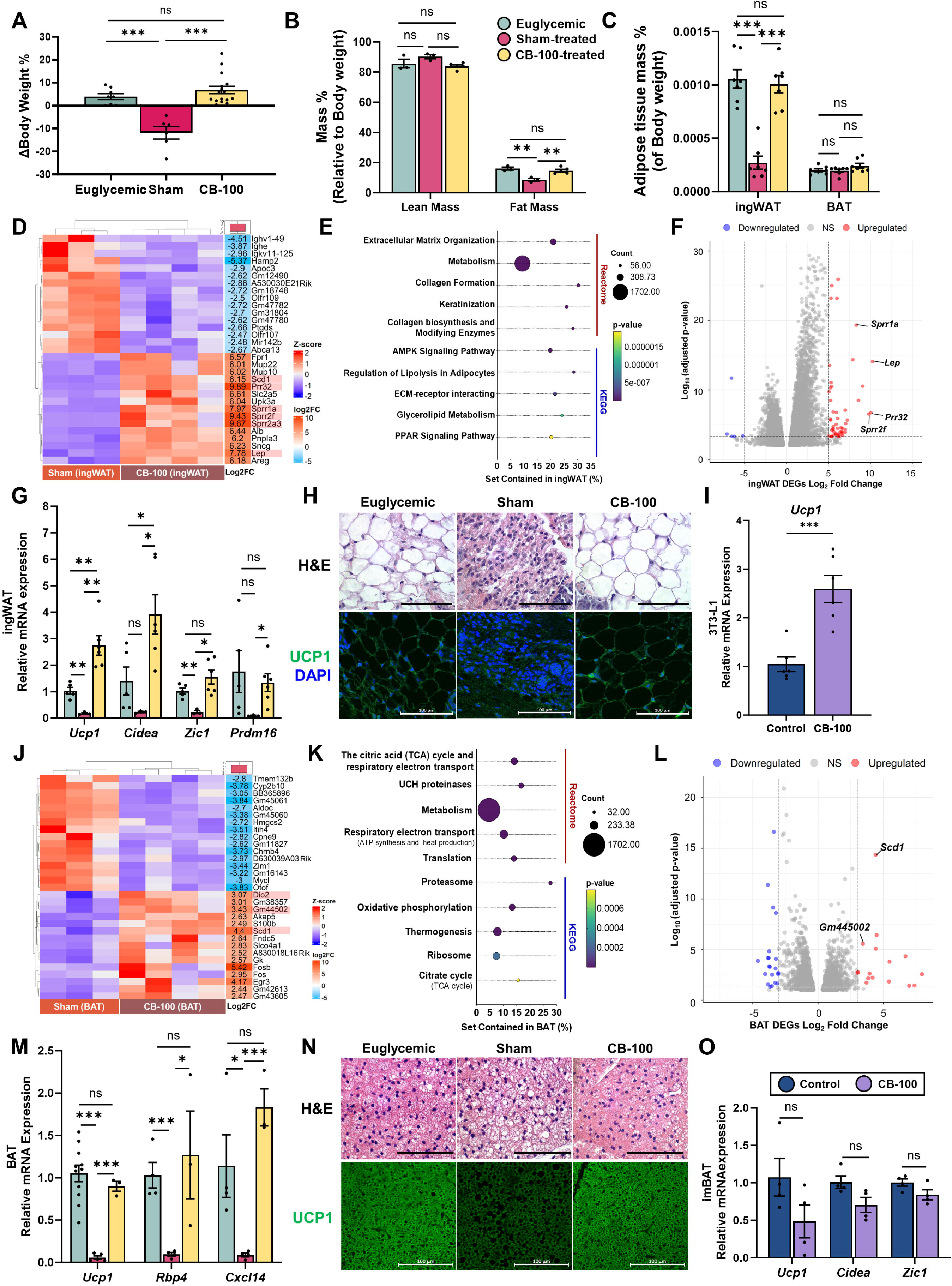
CB-100 Enhances the Adipocyte Differentiation and Browning in White Adipose Tissue and Prevents the Whitening of Brown Adipose Tissue. **(A)** Percentage change of body weight between pre-and post-treatment of euglycemic control (green, n=8), sham-treated (red, n=6), and CB-100 treated NOD mice (yellow, n=16). **(B)** Lean and fat mass percentage relative to body weight of euglycemic (green, n = 3), sham (red, n = 3), and CB-100-treated (yellow, n = 4) female NOD mice. **(C)** Percentage fat mass relative to the body weight of inguinal white adipose tissue (ingWAT) and brown adipose tissue (BAT) from euglycemic (green, n = 7), sham (red, n = 7), and CB-100-treated (yellow, n = 7) female NOD mice. **(D&I)** Cluster analysis of heatmap in sham (n = 3) and CB-100-treated NOD mice (n = 4) for **(D)** inguinal white adipose tissue (ingWAT) and **(I)** brown adipose tissue (BAT) with DEseq2; Top 15 differentially expressed genes (DEGs) with significant upregulation (red) and downregulation (blue) (p_adjust < 0.05, abs (Log_2_FC) > 2, basemean > 50) were represented. Noteworthy genes are highlighted in red. See supplementary tables S4 and S5 for more information. **(E & J)** Top 5 Reactome and KEGG pathway classification from gene-set-based analysis of significantly upregulated DEGs (Log_2_FC > 2, p_val <0.01) in (**E**) ingWAT and (**J**) BAT of CB-100-treated (n=4) compared to sham-treated (n=3) NOD mice. **(F&K**) The volcano plot of DEGs in CB-100-treated NOD mice (n=4) compared to sham-treated (n=3) in **(F)** ingWAT (p_adjust < 0.0005, abs(Log_2_FC) > 5) and **(K)** BAT (p_adjust < 0.05, abs(Log_2_FC) > 3). Upregulated genes are highlighted in red circles, and downregulated genes are highlighted in blue. **(G)** Relative mRNA expression levels of browning marker genes (*Ucp1, Cidea, Zic1,* and *Prdm16*; relative to euglycemic control) in ingWAT from euglycemic control (yellow, n=5), sham-treated (red, n=3), and CB-100-treated (brown, n=6) NOD mice (female). **(H&N)** Representative histology and immunofluorescence images of (**H)** ingWAT and **(N)** BAT from euglycemic control, sham-treated, and CB-100-treated NOD mice with H&E and UCP1 (green) with nuclear staining (DAPI, blue). Scale bar: 100 µm. (**I**) Relative *Ucp1* mRNA expression in control (navy blue) and CB-100 (13.2 μg/mL, 48 hours) treated (purple) 3T3-L1 (white adipocyte) cell line. **(M)** Relative mRNA expression of thermogenic marker genes (*Ucp1, Rbp4*, and *Cxcl14*; relative to euglycemic control) in BAT of euglycemic control (yellow, n=4-10), sham-treated (red, n=4-5) and CB-100 treated (brown, n=3) NOD mice (female). **(O)** Relative mRNA expression of thermogenic marker genes (*Ucp1, Cidea,* and *Zic1*) in control (navy blue) and CB-100 (13.2 μg/mL, 48 hours) treated (purple) imBAT cell line Data are presented as Mean ± SEM. Statistical significance was determined using an unpaired t-test. Groups compared for statistical analysis are indicated by the line. *p < 0.05, **p < 0.01, ***p < 0.001, ns (p > 0.05).

The top 15 upregulated and downregulated differentially expressed genes (DEGs) in CB-100-treated ingWAT versus sham-treated ingWAT reveal a distinct heatmap (Figure 3D). Gene-set-based analyses highlight extracellular matrix organization, metabolism, lipolysis, and glycolipid metabolism (Figure 3E). A volcano plot (p-value < 0.0005, |log2 fold change| > 5) identifies 61 upregulated and 5 downregulated genes (Figure 3F). Among these, genes involved in adipocyte differentiation, maturation, or browning, such as Small Proline-rich Proteins 1A, 2F, and 2A3 (*Sprr1a, Sprr2f, and Sprr2a3*), Proline-rich 32 (*Prr32*), Leptin (*Lep*), and Stearoyl-coenzyme A desaturase 1 (*Scd1*), are notably upregulated (Figure 3D and 3F). Quantitative PCR (qPCR) mRNA expression analysis shows a high expression of browning marker genes (*Ucp1*, *Cidea*, Zic Family Member 1 (*Zic1*) and PR/SET Domain 16 (*Prdm16*)) in ingWAT from CB-100-treated NOD mice compared to sham-treated NOD mice (Figure 3G). Immunofluorescence staining of ingWAT shows increased UCP1 in CB-100-treated NOD mice compared to euglycemic control and sham-treatment (Figure 3H).

To test if CB-100 directly promotes the browning of white adipocytes, the effect of CB-100 on browning was tested *in vitro*. 3T3-L1 cells treated with CB-100 for two days demonstrate increased *Ucp1* mRNA expression (Figure 3I). These results collectively indicate that CB-100 treatment promotes adipocyte differentiation and the browning of white adipose tissue, which likely contributes to enhanced glucose homeostasis.

### CB-100 Prevents the Whitening of Brown Adipose Tissue

BAT is recognized for its role in glucose homeostasis through increased energy expenditure and enhanced insulin sensitivity ^20^. In addition to the reported BAT whitening in the context of obesity and type 2 diabetes ^21^, we observe a reduction in BAT thermogenic activity marker genes (*Ucp1*, Retinol binding protein 4 (*Rbp4*), and Chemokine (C-X-C motif) ligand 14 (*Cxcl14*)) with the development of T1D symptoms in NOD mice (Figure 3M).

Although BAT mass is similar between CB-100-treated and sham-treated NOD mice (Figure 3C), the DEGs heatmap reveals distinct expression patterns in endogenous BAT of CB-100-treated compared to sham-treated NOD mice (Figure 3J). Gene-set-based Reactome and KEGG pathway analyses of CB-100-treated BAT compared to sham-treated reveal enriched metabolism and thermogenic activity pathways (Figure 3K). A volcano plot (p-value < 0.05, |log_2_ fold change| > 3) shows 16 significantly upregulated and 22 downregulated genes (Figure 3L). Notably, genes associated with BAT thermogenesis, such as iodothyronine deiodinase 2 (*Dio2*), stearoyl-coenzyme A desaturase 1 (*Scd1*), and the long non-coding RNA Gm445002, are upregulated (Figures 3J and 3L). DEGs analysis indicates enhanced thermogenic activity in BAT of CB-100-treated NOD mice compared to sham (Figures 3I-3K). However, there are no significant differences in the expression levels of thermogenesis marker genes (*Ucp1, Rbp4*, and *Cxcl14*) between euglycemic control and CB-100-treated NOD mice (Figure 3M). Consistent with the *in vivo* data, *in vitro* experiments with imBAT treated with CB-100 show no significant changes in thermogenesis-related gene expressions (*Ucp1, Cidea,* and *Zic1*) compared to untreated (Figure 3O).

H&E staining of BAT showed enlarged lipid droplets in sham-treated NOD mice compared to euglycemic control and CB-100 treated NOD mice, while there is no noticeable difference in lipid droplets between euglycemic control and CB-100 treated NOD mice (Figure 3N). Immunofluorescence staining of BAT from CB-100-treated NOD mice reveals a higher UCP1 expression compared to the sham-treated NOD mice, but no significant differences were observed compared to non-diabetic control mice (Figure 3N). These findings suggest that CB-100 treatment prevents the reduction of BAT thermogenic activity due to the development of T1D symptoms in NOD mice.

### CB-100 Enhances Glucose Uptake in Skeletal Muscle, Liver, and Adipose Tissue

Immunofluorescence of gastrocnemius muscles from non-diabetic, CB-100-treated, and sham-treated NOD mice reveals higher colocalization of GLUT4 and α-sarcoglycan in CB-100-treated mice (Figures 4A and 4B). This finding suggests enhanced GLUT4 translocation to the plasma membrane, facilitating increased glucose uptake. In the liver, successful CB-100-treated mice show higher liver glycogen level than the sham and failed treatment (Figure 4C).

**Fig 4.**
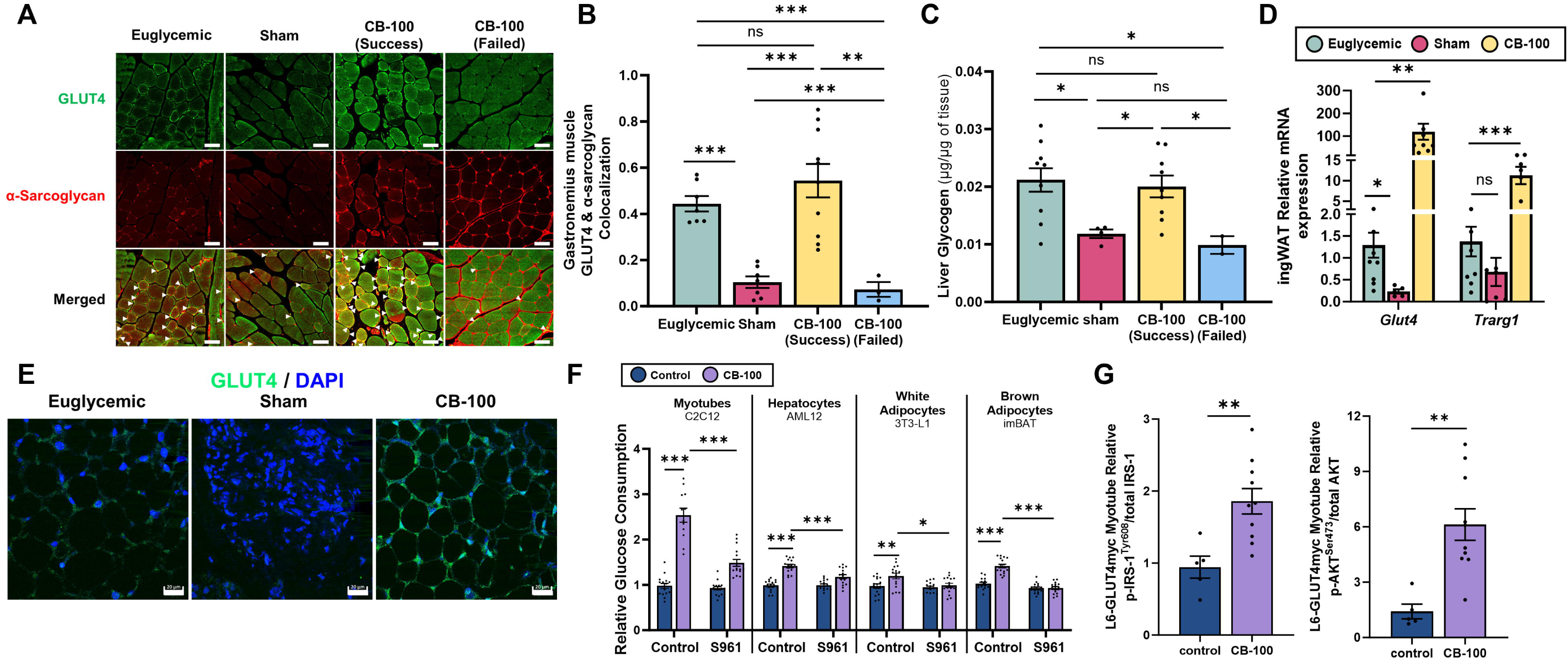
CB-100 Enhances Glucose Uptake in Skeletal Muscle, Liver, and Adipose Tissue. (**A**) Representative immunofluorescence images of the colocalization of GLUT4 (green) with α-sarcoglycan (a plasma membrane marker, red) in gastrocnemius muscle tissues of euglycemic control, sham-treated, and CB-100 treated (success and failed) NOD mice. Colocalizations are denoted by arrows Scale bar: 50 μm. (**B**) Quantitative analysis of GLUT4 and α-sarcoglycan colocalization in euglycemic control (green, n=3), sham-treated (red, n=3), and CB-100-treated (successful; yellow, n=5 & failed; blue, n=2) NOD mice. Arrows denote colocalizations. Scale bar: 50 μm. (C) Hepatic glycogen levels measured in euglycemic control (green, n=10), sham-treated (red, n=4), and CB-100-treated (successful; yellow, n=9 & failed; blue, n=2) NOD mice, normalized to tissue weight. (D) Relative expression levels of insulin-related glucose transporter marker genes (Glucose Transporter Type 4 (*Glut4*) and Trafficking Regulator of GLUT4 1 (*Trarg1*)) in ingWAT of euglycemic (green n=9), sham-treated (red, n=5) and CB-100-treated (yellow, n=9) NOD mice (female). (**E**) Immunofluorescence of GLUT4 (green) and nuclei (blue) in inguinal white adipose tissue (ingWAT) of euglycemic control, sham-treated, and CB-100 treated NOD mice. Scale bar: 20 μm. (**F**) Relative glucose consumption in myotubes (C2C12), hepatocytes (AML12), white adipocytes (3T3-L1), and brown adipocytes (imBAT) treated without (control, navy blue) or with CB-100 (purple, 52.8 µg/mL) and in the absence (control) or presence of S961 (200 nM) for 24 hours. (**G**) Immunoblot analysis of p-IRS-1^Tyr606^/total-IRS-1 and p-AKT^Ser473^/total-AKT from L6.GLUT4Myc myotube untreated (navy blue, control) or treated with CB-100 (purple, 10μg/mL, 16∼18 hours) then spiked in insulin (10nM, 5minutes). Data are presented as mean ± SEM. Statistical significance was assessed using an unpaired t-test, with significance denoted as follows: *p < 0.05, **p < 0.01, ***p < 0.001, ns (p > 0.05). The groups compared for statistical analysis are connected by lines.

Immunofluorescence of ingWAT from non-diabetic, CB-100-treated, and sham-treated NOD mice reveals increased expression of GLUT4 in CB-100-treated NOD’s adipose tissue compared to sham-treated (Figure 4E). Moreover, relative gene expression in ingWAT shows upregulation of insulin-regulated glucose transporter trafficking-related genes (Glucose transporter type 4 (*Glut4*) and Trafficking regulator of GLUT4 1 (*Trarg1*) ^22^) in CB-100-treated NOD mice compared to the sham treatment (Figure 4D). On the other hand, BAT does not show a high expression of GLUT4 as BAT in euglycemic control mice (Figures S2K and S2L). *In vitro* analyses of myotubes, hepatocytes, and adipocytes show enhanced glucose uptake with CB-100 and diminished effects following exposure to the insulin receptor antagonist S961 (Figure 4F). Western blot analysis shows increased phosphorylation of insulin receptor substrate 1 (IRS-1) and protein kinase B (AKT) in L6-GLUT4myc myotubes upon treatment with CB-100 (Figures 4G and S3A). These findings collectively suggest that CB-100 treatment enhances glucose uptake in adipose tissue, skeletal muscle, and liver, at least partially through the insulin receptor pathway, which leads to improved glucose homeostasis.

### Identification of Nidogen-2 as a Brown Adipocyte-Secreted Glucagon Secretion Regulator Protein

We utilized a series of separation techniques to identify the glucagon secretion-inhibiting proteins underlying the effects of CB-100. The CB-100 fraction was subjected to anion-exchange and size-exclusion chromatography (SEC), and fractions from SEC were evaluated by mass spectrometry and correlated with their ability to inhibit glucagon secretion (Figure 5A). Mass spectrometry correlation analysis and Pearson’s correlation coefficient versus protein rank show the top candidate secreted proteins (Figure 5B), which we tested for glucagon inhibition in various concentrations (Figures S4A, S4C, and S4E). Among the candidate proteins, nidogen-2 is strongly associated with glucagon secretion inhibition (Figure 5C). Based on its robust and consistent inhibition of glucagon secretion from murine and human islets, we further investigated nidogen-2. Immunofluorescence shows intracellular nidogen-2 in fully differentiated imBAT (Figure 5D), and western blot analysis confirms nidogen-2 in the conditioned buffer from both eBAT and imBAT (Figure 5E). However, the cellular secretions show a cleaved nidogen-2 fragments rather than only the full-length protein. Based on the elution volumes of SEC fractions with nidogen-2 detected (Figure 5C) and SEC calibration curve, the molecular sizes of 3 peaks (Figure 5C) were calculated to be 84-109kDa, 57-90kDa, and 23-34kDa (Figure 5F).

**Figure 5.**
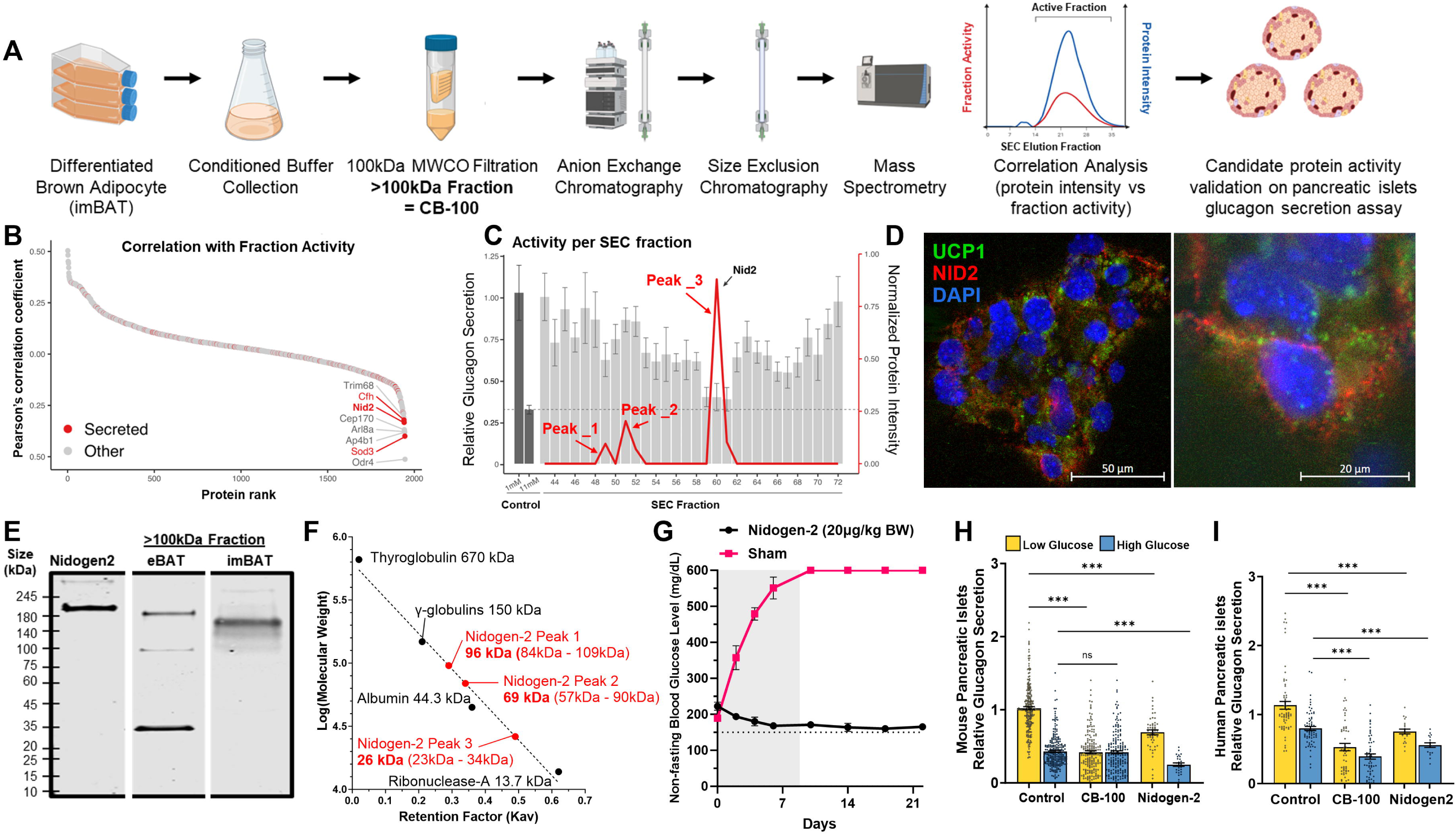
Identification of Nidogen-2 as a Brown Adipocyte-Secreted Glucagon Secretion Regulator Protein. (**A**) Schematic of the experimental design and process for identifying active proteins from CB-100. (**B**) Rank order of protein signal intensity correlation in size exclusion chromatography fractions with the glucagon secretion inhibition activity. Reported secreted proteins are indicated in red circles. (C) Plot of glucagon secretion level with size exclusion chromatography fractions at low glucose (1mM) condition (grey bar) and nidogen-2 protein intensity (red). (For glucagon secretion, the control condition (1mM and 11mM) is indicated with the left two dark grey bars. The expected lowest glucagon secretion is determined with the glucagon secretion level at high glucose condition (11mM) and indicated with the dotted line. (D) Immunofluorescence image of nidogen-2 (red), UCP1 (green), and nucleus (blue) in fully differentiated brown adipocytes (imBAT) Scale bar: 50 μm (left) and 20 μm (right). (**E**) Western blot analysis of nidogen-2 in recombinant nidogen-2 (Leu31-Lys1403) and >100kDa conditioned buffer from embryonic brown adipose tissue (eBAT) and immortalized brown adipocytes (imBAT). (**F**) Interpolated molecular weight of nidogen-2 peaks in SEC elution shown in figure 5C using linear regression from log (molecular weight) versus retention factor (K_av_) of molecular weight standard mix which includes thyroglobulin (670kDa), γ-globulins (150kDa), albumin (44.3kDa), and ribonuclease-A (13.7kDa). (**G**) Non-fasting blood glucose levels measured following 7 days of injection of recombinant mouse nidogen-2 (Leu31-Lys1403) (black, subcutaneous, 20μg/kg BW, n = 5) and sham (red, KRBH, n = 3) in NOD mice (female, 18-22 weeks old). Normal non-fasting blood glucose level for NOD mice was indicated with a dotted line (150mg/dL). Grey highlighted days are the days of the nidogen-2 injections. **(H&I)** Relative glucagon secretion measured in low (yellow, 1mM) and high (teal, 11mM) glucose conditions in non-diabetic **(H**) mouse (n=15) and (**I**) human (n=4) pancreatic islets without or with CB-100 (2.5μg/mL) or recombinant mouse nidogen-2 (Leu31-Lys1403) (40ng/mL). Data are presented as Mean ± SEM. Statistical significance was determined using an unpaired t-test. Groups compared for statistical analysis are indicated by the line. *p < 0.05, **p < 0.01, ***p < 0.001, ns (p > 0.05).

When recombinant mouse nidogen-2 (Leu31-Lys1403) was injected into early-stage hyperglycemia NOD mice, animals showed recovery of euglycemia (Figure 5G). Moreover, recombinant mouse nidogen-2 suppresses glucagon secretion in mouse and human pancreatic islets (Figures 5H and 5I). These findings identify nidogen-2 as a brown adipocyte-secreted regulatory protein for hyperglycemia and glucagon secretion.

### Regulation of Intracellular Messengers of Glucagon Secretion by CB-100 and Nidogen-2 in Pancreatic α-cells

To investigate the mechanisms underlying glucagon secretion inhibition by CB-100 and nidogen-2, we examined α-cell signaling pathways. Both CB-100 and nidogen-2 inhibit glucagon secretion in dispersed islet cells from mouse and human islets under low and high glucose conditions (Figure 6A). These data indicate that CB-100 and nidogen-2 influence α-cells directly rather than indirectly through other islet cell types. To determine if specific receptors on pancreatic α-cells mediate the inhibition, we tested antagonists for the glucagon receptor (GCGR), GLP-1 receptor (GLP1R), somatostatin receptor (SSTR), EphA4 receptor (EphA4R), and insulin receptor (INSR) on CB-100 or nidogen-2 activity. We observe no changes in these activities after antagonizing GCGR, GLP-1R, SSTR, or EphA4R (Figure S1D). However, the INSR antagonist (S961) diminishes the inhibitory effects of CB-100 and nidogen-2 under low and high glucose conditions (Figure 6B).

**Figure 6.**
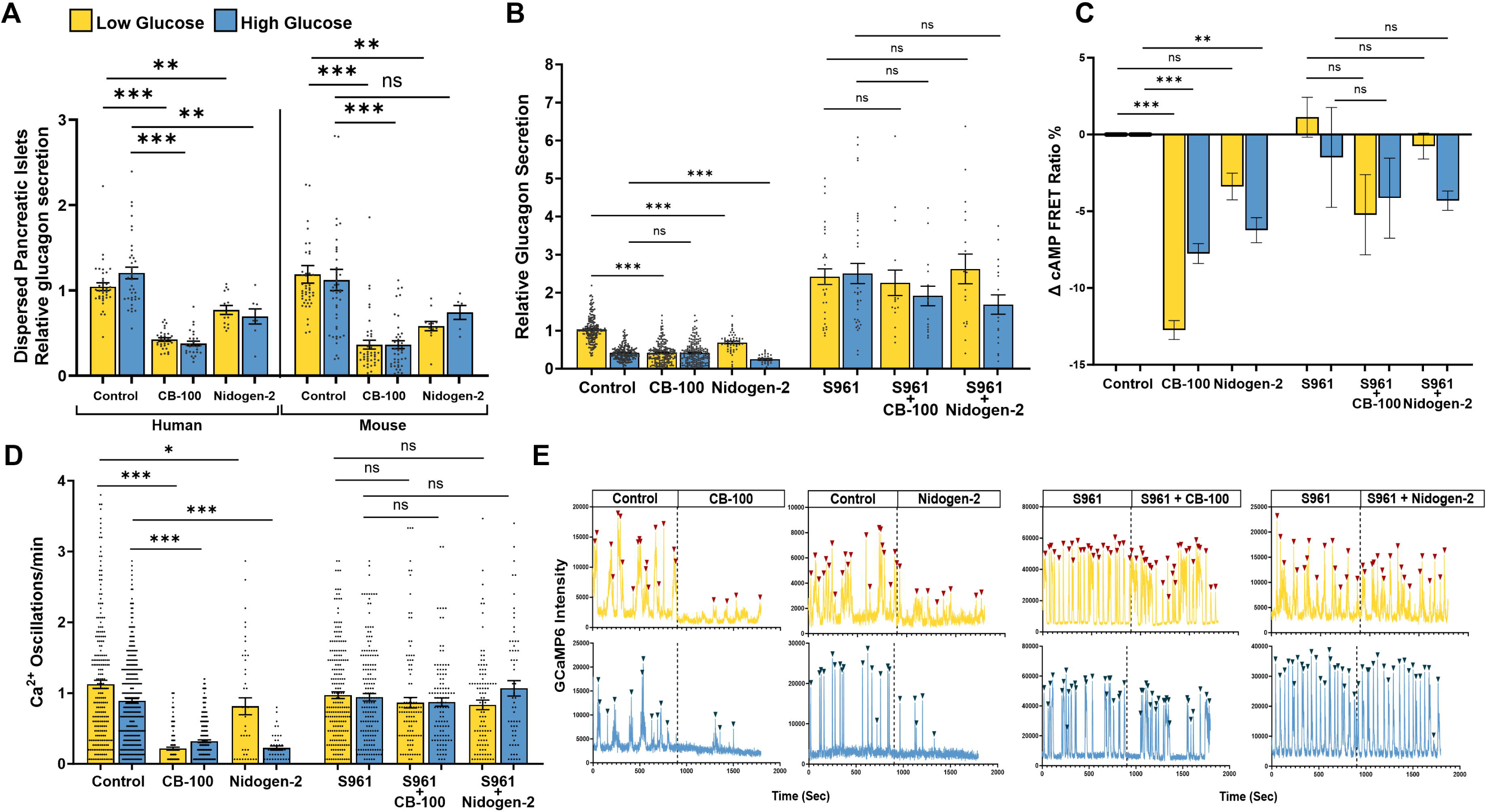
Regulation of Intracellular Messengers of Glucagon Secretion by CB-100 and Nidogen-2 in Pancreatic α-cells. (**A**) Relative glucagon secretion measured in low (yellow, 1mM) and high (teal, 11mM) glucose conditions in non-diabetic mouse (n=7) and human (n=4) dispersed pancreatic islets without or with CB-100 (2μg/mL) and nidogen-2 (2μg/mL for human, 2ng/mL for mouse). (**B**) Relative glucagon secretion measured in low (yellow, 1mM) and high (teal, 11mM) glucose conditions with or without S961 (1 μM) and CB-100 (2 μg/mL) or Nidogen-2 (40 ng/mL) in mouse (non-diabetic, n=8) pancreatic islets. (C) Percentage change calculated from cAMP FRET intensity ratio in non-diabetic mouse pancreatic islets (n=5) in low (yellow, 1 mM) and high (teal, 11 mM) glucose conditions before and after treatment (CB-100 (5 μg/mL) or nidogen-2 (40 ng/mL)) with or without S961 (1 μM). (D) Ca^2+^ oscillation frequency measured in low (yellow, 1mM) and high (teal, 11mM) glucose conditions in non-diabetic mice (n=7) with or without CB-100 (5 μg/mL) or Nidogen-2 (40 ng/mL) and S961 (1 μM). (**E**) Representation of calcium indicator (GCaMP6) fluorescence intensity tracing in individual mouse pancreatic α-cells in low (yellow, 1mM) and high (teal, 11mM) glucose conditions with CB-100 (5 μg/mL) or Nidogen-2 (40 ng/mL) and with or without S961 (1 μM). Data are presented as Mean ± SEM. Statistical significance was determined using an unpaired t-test. Groups compared for statistical analysis are indicated by the line. *p < 0.05, **p < 0.01, ***p < 0.001, ns (p > 0.05).

We next measured intracellular cAMP and Ca^2+^ activity by live cell imaging of α-cells in mouse islets. CB-100 and nidogen-2 inhibit both intracellular cAMP and Ca^2+^ activity compared to control (Figures 6C-6E). These inhibitions of intracellular cAMP and Ca^2+^ activities are blocked by the insulin receptor antagonist, S961 (Figures 6C-6E). These results indicate that the suppression of glucagon secretion by CB-100 and nidogen-2 in pancreatic α-cells is mediated, at least in part, through an insulin receptor-dependent manner.

## DISCUSSION

We investigated the mechanisms underlying the restoration of euglycemia following embryonic BAT transplant into T1D mouse models. Our findings suggest that the hormone action of protein(s) in the large (>100kDa) BAT-secreted fraction (CB-100) mediates positive metabolic effects. Similar to the effects observed with embryonic BAT transplantation, subcutaneous injection of CB-100 maintains sustained euglycemia (Figure 2B), with a strong association between plasma glucagon levels and treatment success. Compared to sham and failed CB-100 treatments, successfully treated NOD mice exhibit near-normal plasma glucagon levels and pancreatic glucagon content (Figure 2F and 2H), suggesting that CB-100 suppresses hyperglucagonemia and α-cell hyperplasia. Early-stage treatment of hyperglycemia with CB-100, when detectable β-cells are still present, significantly improves the success rate (Figures S2A and S2D). Immunofluorescence analysis of NOD mice supports this mechanism since pancreata from successful CB-100 treated animals have some detectable insulin-positive β-cells (Figure 2J). Although partial restoration of plasma insulin is observed in successfully treated mice, these levels remain significantly lower than those in euglycemic control animals. The presence of insulin positive β-cells and plasma insulin could be attributed to the restoration of euglycemia in CB-100 treated mice. Chronic hyperglycemia is reported to exacerbate reactive oxygen species production and glucotoxicity, and heighten insulin resistance necessitates increased insulin demand from β-cells ^23^. Therefore, effective blood glucose control and improve insulin sensitivity via CB-100 could alleviate β-cell stress, thereby enhancing their functionality and survival. This improvement is reflected in the increased pancreatic and plasma insulin levels and the presence of insulin-positive β-cells in CB-100-treated mice (Figures 2G, 2I, and 2J). However, it is important to note that plasma insulin levels and pancreatic insulin content do not correlate with the efficacy of CB-100 treatment (Figures 2G and 2I). Instead, plasma glucagon levels are indicative of treatment success (Figures 2F and 2H), aligning with prior studies that highlight the therapeutic benefits of suppressing hyperglucagonemia in type 1 diabetes^14,24^.

Within CB-100, we identified nidogen-2 as playing a central role in reversing hyperglycemia in a T1D model (Figure 5F). Recombinant nidogen-2 suppresses glucagon secretion from pancreatic islets (Figures 5G and 5H) and enhances glucose uptake in adipocytes (Figure S4F). While full-length nidogen-2 suppresses glucagon secretion, its suppressive effect is not as strong as that observed with CB-100 (Figures 5G and 5H). Both CB-100 and nidogen-2 suppress glucagon secretion from dispersed islet cells, indicating a direct modulation of α-cell function (Figure 6A). Live cell imaging shows that both CB-100 and nidogen-2 decrease intracellular cAMP and Ca^2+^ activity, and that these effects are attenuated by antagonizing the insulin receptor (Figures 6B-6E). CB-100 and nidogen-2 increase glucose uptake in brown adipocytes *in vitro* (Figures 4F and S4F), while CB-100 also increases glucose uptake in myotubes and hepatocytes (Figure 4F). This stimulatory effects on glucose uptake are mitigated when insulin receptors are blocked (Figures 4F and S4G). CB-100 improves *in vivo* glucose uptake in adipose tissue, skeletal muscle, and liver (Figures 4A-E). Upregulation of GLUT4 in ingWAT (Figures 4D and 4E) and increased phosphorylation of insulin signaling pathway components (IRS-1 and AKT) ^25^ (Figure 4G) by CB-100 further implicates an important role of the insulin receptor in CB-100-mediated glucose uptake. Moreover, the effects of CB-100 and nidogen-2 are reduced by insulin receptor antagonism. Taken together, these results suggest that nidogen-2 plays a major role in the action of CB-100, and the activity of nidogen-2 is mediated, at least in part, by the insulin receptor.

Nidogen-2 has been reported as an extracellular matrix protein involved in embryo development, basement membrane assembly, tissue regeneration, and interaction with other extracellular matrix proteins ^26–30^. Our data show that in many respects, treatment with recombinant full-length nidogen-2 mimics previous embryonic BAT transplant data ^13,14^ and replicates the actions of CB-100 (Figures 5G-5I and Figure 6). By other measures, its activity is less pronounced or not observed at the concentration tested (Figures 5H, 5I, and S4F). However, we detect cleavage products in CB-100 and eBAT secretions that are smaller than the reported size of nidogen-2 (200kDa) (Figure 5E) ^31^. SEC fractions with nidogen-2 detected show three major peaks (Figure 5C) with one overlapping the fraction of highest activity. The estimated size of all three peaks combined is 164-233kDa (Figure 5F), which encompasses the reported molecular weight of full-length nidogen-2. These results suggest that a cleavage product(s) of nidogen-2 may be necessary to achieve its maximum mediate metabolic effects.

BAT loses thermogenic activity and insulin sensitivity during obesity and type 2 diabetes ^20^. Our data is consistent with this observation by showing BAT “whitening,” which is indicated by decreased thermogenic marker gene expression with T1D phenotype progression in NOD mice (Figure 3L). Compared to sham-treated hyperglycemic NOD mice, DEGs analysis of endogenous in BAT following CB-100 treatment reveals upregulation of *Dio2*,*Gm445002,* and *Scd1*, genes to promote adipogenesis, adipocyte browning, and thermogenesis (Figures 3I and 3K) ^32–35^. Moreover, gene-set based analysis highlighted metabolism and thermogenesis as enriched pathways (Figure 3J), which implies the increase metabolic and thermogenic activity of BAT in CB-100 treated NOD mice. However, further qPCR mRNA expression does not demonstrate a notable difference in expression of thermogenesis marker genes (*Ucp1, Rbp4,* and *Cxcl14*) between CB-100 treated and euglycemic controls (Figure 3L), suggesting that CB-100 prevents the whitening of BAT instead of further improving its thermogenic activity.

In addition to preventing BAT whitening, our results show that CB-100 promotes differentiation and browning of white adipose tissue, thereby facilitating better glucose utilization and energy expenditure. Metabolic studies of live animals show higher oxygen consumption (VO_2_, VCO_2_/VO_2_) in CB-100-treated compared to sham-treated animals (Figures 2C and S2G), while there is no significant difference in physical activity or heat expenditure (Figures S2H and S2I). The increased glucose utilization per energy expenditure is reflected in the maintenance of body weight and fat mass (Figures 3A and 3B). Mice treated with CB-100 (RER close to 1.00) utilize glucose as their major energy source, while sham-treated mice (RER close to 0.70) utilize fat instead ^36^ (Figure 2C). By increasing efficient glucose utilization and preserving function of adipose tissue, CB-100-treated NOD mice appear to enhance glycemic control by mediating glucose disposal and secreting beneficial adipokines ^37^. H&E staining of ingWAT in sham-treated NOD mice shows fibrotic tissue that constrains adipocyte expansion (Figure 3N), which is expected to promote ectopic fat deposition and result in fat accumulation in other organs like the liver ^38^. Supporting this concept, liver samples from sham-treated NOD mice display small-droplet macro-vesicular steatosis (sd-MaS) fat accumulation, while CB-100-treated and euglycemic NOD mice’ livers do not (Figure S2J).

DEGs analysis of ingWAT from successful CB-100 treated animals highlights the upregulation of several proline-related genes (*Sprr1a, Sprr2f, Sprr2a3*, and *Prr32*), *Lep,* and *Scd1* (Figures 3D and 3F). Proline serves as a primary precursor to extracellular collagen, a major component of the extracellular matrix (ECM) ^39^. Basement membrane (BM)-associated collagens support adipose tissue differentiation and preadipocyte maturation, and ECM molecules are involved in adipose function regulation, such as energy metabolism ^40^. Thus, increased proline levels within ingWAT could underscore the role of CB-100 in enhancing adipocyte functionality ^39^. In addition to its actions on glucagon secretion, nidogen-2 within CB-100 may contribute to adipocyte proliferation by acting as a linker molecule ^41^ between basement membrane proteins, thereby supporting the growth and turnover of ingWAT. The observed upregulation of *Lep* suggests that the CB-100 benefits may be related to the regulation of food intake, increasing energy expenditure, and browning in ingWAT ^42,43^. Similarly, upregulation of *Scd1* is associated with promoting adipogenesis and browning of adipocytes in ingWAT^33,44^. Consistent with ingWAT browning, we observe comparable or higher expression of the thermogenic marker (*Ucp1, Cidea, Zic1*, and *Prdm16*) and protein (UCP1) in ingWAT from CB-100-treated NOD mice compared to non-diabetic NOD mice (Figures 3G and 3N). Finally, the gene-set-based analysis highlights metabolism, glycerolipid metabolism, and lipolysis-related pathways (Figure 3E). Glycerolipid (GL) metabolism and lipolysis result in the production of free fatty acids (FFA)^45^. GL/FFA cycling has been reported to improve energy homeostasis and thermogenesis through the peroxisomal proliferator-activated receptor (PPAR) and AMP-activated protein kinase (AMPK) signaling pathways^45^. GL/FFA from adipose tissue has been suggested to be involved in regulating whole-body glucose homeostasis and insulin sensitivity ^46,47^. These results collectively suggest that CB-100 treatment not only promotes the recovery of white adipose tissue mass but also induces browning, thereby enhancing energy expenditure and glucose management.

Overall, this study reports establishes a novel hormonal mechanism that underlies the reversal of diabetes phenotypes in T1D mouse models by embryonic BAT transplants. We show a critical role of large (>100kDa) BAT-secreted proteins in promoting glucose homeostasis by regulating glucagon secretion, activating the insulin receptor, preventing the whitening of BAT, and promoting the browning of ingWAT. We identify a key protein, nidogen-2, which mimics many important actions of the entire CB-100 fraction. The broad metabolic effects of CB-100 and nidogen-2 are detailed through *in vitro* and *in vivo* assessments, unveiling a therapeutic strategy for diabetes management beyond insulin-centric paradigms. Future research on identifying and characterizing nidogen-2 cleavage products as well as other possible active components within CB-100 will elucidate their roles in glycemic and metabolic regulation.

## LIMITATION OF STUDY

Our study has provided valuable insights into the role of CB-100 and nidogen-2 in glucose homeostasis, yet we recognize several limitations. CB-100 is a complex mixture comprising various proteins, which complicates pinpointing any specific protein responsible for the observed metabolic effects. Although nidogen-2 mimics many of the effects seen with CB-100, it does not account for all effects, and we do not know whether its cleavage products or other CB-100 components play participate in the overall mechanism. We have identified insulin receptor-dependent pathways as crucial for glucagon regulation, but other potentially involved pathways remain to be fully elucidated. Our research primarily addresses the immediate metabolic effects without exploring the long-term consequences on systemic physiology and other metabolic pathways. This limitation highlights the necessity for ongoing research to unravel the intricate signaling networks at play and comprehensively evaluate the full therapeutic potential of BAT-secreted proteins.

## ACKNOWLEDGEMENT

This work was supported by the US National Institutes of Health (grants R01DK123301 and R01DK115972) and The Leona M. and Harry B. Helmsley Charitable Trust (grants G-2305-06050, G-1912-03558, and G-2001-04215). Human pancreatic islets and/or other resources were provided by the NIDDK-funded Integrated Islet Distribution Program (IIDP) (RRID:SCR _014387) at City of Hope, NIH Grant U24DK098085 and the JDRF-funded IIDP Islet Award Initiative. T1D donor islet isolation was also supported by The Leona M. and Harry B. Helmsley Charitable Trust (G-2018PG-T1D027). For the *in vivo* metabolic studies, we thank the Diabetes Models Phenotyping Core at Washington University School of Medicine. NIH grant P30 DK020579. Live cell and immunofluorescence imaging were performed with a Zeiss LSM 880 at the Washington University Center for Cellular Imaging (WUCCI) supported by Washington University School of Medicine, Diabetes Research Center (P30DK020579), The Children’s Discovery Institute of Washington University and St. Louis Children’s Hospital (CDI-CORE-2015-505 and CDI-CORE-2019-813) and the Foundation for Barnes-Jewish Hospital (3770 and 4642). For RNA sequencing, we thank the Genome Technology Access Center (GTAC) at the McDonnell Genome Institute at Washington University School of Medicine. GTAC is partially supported by NCI Cancer Center Support Grant P30 CA91842 to the Siteman Cancer Center. This publication is solely the responsibility of the authors and does not necessarily represent the official view of NCRR or NIH. Experiment with L6-GLUT4myc myotube cell lines were supported by R01DK102233 to DCT; AHA fellowship (#905770). We thank the laboratory of Rajan Sah at Washington University in St. Louis for providing a C2C12 myotube cell line. For schematic figures (Figure 2A and 5A) BioRender.com was used.

## Author contributions

J.L. designed, performed, analyzed overall experiments and data, and wrote the manuscript. A.U. established the CB-100 anion exchange chromatography and size exclusion chromatography fractionation system described in Figures 5A and 5F. D.G and E.M.W. performed mass spectrometry, proteomic analysis, and correlation analysis described in Figures 5A-C, 5J, S4B and S4D. R.B. and D.T. performed and analyzed the western blot analysis with L6-GLUT4myc myotube described in Figure 4G, S3A, S3C and S3D. D.W.P. conceptualized the project and designed experimental approaches. All authors participated in editing the manuscript.

## STAR METHODS

### Resource availability Lead contact

Further information and requests for resources and reagents should be directed to and will be fulfilled by the lead contact, Jeongmin Lee.

### Materials availability

This study did not generate new unique reagents.

### Data and code availability

-All of the data supporting this study are included in the article. All raw data used to generate the figures throughout the manuscript can be found within the Data S1 document.
-Raw RNA-seq data have been deposited at GEO and are publicly available as of the date of publication. Accession numbers are listed in the key resources table.
-Mass spectrometry data have been deposited at ProteomeXchange and are publicly available as of the date of publication. PXD are listed in the key resources table.
-Microscopy data reported in this paper will be shared by the lead contact upon request.
-This paper does not report original code.
-Any additional information required to reanalyze the data reported in this paper is available from the lead contact upon request.

## EXPERIMENTAL MODELS AND SUBJECT DETAILS

### Cell culture

Immortalized brown preadipocyte (imBAT, derived from stromal vascular fraction of brown adipose tissue) cells were provided by M. Christian, Ph.D. (University of Warwick, Division of Metabolic and Vascular Health ^48^). Preadipocyte differentiation was induced by culturing with differentiation media (DMEM/F12 + GlutaMAX, 10% (v/v) fetal bovine serum (FBS), 1% (v/v) Penicillin-streptomycin (Pen/Strep), 170nM insulin, 250nM Dexamethasone, 0.5mM IBMX, 0.1nM T3, 125uM Indomethacin) at 33°C for 48 hours. Differentiated imBAT cells were then cultured with maintenance media (DMEM/F12 + GlutaMAX, 10% (v/v) FBS, 1% (v/v) Pen/Strep, 170nM insulin, 0.1nM T3) at 37°C for 8 days, as previously described. ^48–51^.

Myotube C2C12 cells were cultured with DMEM, 20% FBS, 1% Penicillin-streptomycin to 60-70% confluency. Once they reached the desired confluency, cells were differentiated (DMEM, 2% (v/v) Horse serum, 1% (v/v) Pen/Strep, change media every 2-3 days) for 10 days before being ready for the experiment as previously described ^52^.

3T3-L1 adipocytes (ATCC) were differentiated (DMEM, 10% (v/v) Bovine Calf Serum (BCS), 1% (v/v) Pen/Strep, 0.5mM IBMX, 1uM Dexamethasone, 170nM Insulin) and maintained (DMEM, 10% (v/v) BCS, 1% (v/v) Pen/Strep, 170nM Insulin) as described ^53^.

Rat L6-GLUT4myc skeletal muscle cells expressing c-myc-tagged GLUT4 protein (Kerafast) were grown as monolayers in MEM-α medium supplemented with 10% (v/v) FBS and 1% (v/v) antibiotic–antimycotic solution (Thermo Fisher). L6-GLUT4myc myoblasts were differentiated into myotubes by incubating in MEM-α medium containing 2% TBS and 1% antibiotic–antimycotic solution. For western blot analysis, L6-GLUT4myc myotubes were pre-treated with CB-100 (10μg/mL) for 16 to 18 hours followed by 1 hour of serum starvation before insulin stimulation (10nM) for 5 minutes prior to isolating protein.

### Experimental models

All mouse experiments were performed under approval of the Washington University Institutional Animal Care and Use Committee. 8–14-week-old, male and female wild-type C57BL/6 (Jackson Laboratory) were used for the islet isolation. For pancreatic α-cell calcium imaging, mice with α-cells expressing GCaMP6f (GCG-iCre-GCaMP6 mice) have been previously described ^54^. For cAMP measurement in α-cells, mice expressing red fluorescent protein in pancreatic α-cells (αRFP mice; GCGCre-RFP) have been previously described ^55^. For the CB-100 injection experiment, NOD/ShiLtJ (Jackson Laboratory) mice (Male and female) were ordered at 9-weeks-old, and blood glucose level was checked weekly to monitor the status of the development of diabetes. Both male and female NOD mice eventually developed hyperglycemia. However, the time and the rate of conversion were notably longer and lower in male mice compared to females. To ensure a consistent and reliable experimental timeline, after the initial two rounds of injections, we opted to conduct the remaining repetitive experiments required for in-depth *in vivo* characterization exclusively with female NOD mice.

Non-diabetic and Type 2 diabetic human islets were received from the Integrated Islet Distribution Program (IIDP), and type 1 diabetic human islets were received from Imagine Pharma (Pittsburgh, PA) through the Helmsley Trust T1D donor islet program. Received islets were cultured in islet media overnight before use. Islet donor information is available in Supplementary Table S1.

## METHOD DETAILS

### Pancreatic islet isolation and culture

Murine pancreatic islets were isolated by digesting the pancreas from C57BL/6 mice (10-14 weeks old) with 12mg Collagenase P (Roche) at room temperature with rotation for 35 minutes. After digestion, islets were washed twice with 10mL of G-Solution (HBSS with 1% BSA) by centrifuging at 300g for 3 minutes and discarding the supernatant. After washing, pellets containing islets were manually picked or isolated through density gradient centrifugation.

For manual picking, the pellet was reconstituted into islet media (RPMI 1640 with 10% FBS, 1% Penicillin-streptomycin, 20 mM HEPES, 11 mM glucose) and placed into petri dishes. Under a dissection microscope, islets were picked using a 10µL pipette and placed in a new dish containing 10mL of islet media for overnight recovery at 37°C. For density gradient islets isolation, the pellet was reconstituted into 10mL of G-Solution and filtered into a 50mL conical tube through a metal strainer (500μm), and an additional 10mL of G-Solution was added to the filtrate. Islets were pelleted by centrifuge at 300g for 3 minutes, and the supernatant was removed. The pellet was resuspended in 15mL of histopaque-1100 (45.45% Histopaque 1077, 54.54% Histopaque 1119, room temperature) and centrifuged at 290g for 20 minutes. The supernatant was transferred to a new 50mL tube with 25mL of G-solution. Islets were pelleted by centrifuging at 450g for 5 minutes, and the supernatant was removed. Islet pellet was washed with 10mL of G-solution by centrifuging at 300g for 3 minutes. The washed pellet was resuspended to 5mL of islet media and transferred to a petri dish with 5mL of islet media. Islets were incubated at 37°C overnight for recovery before the experiment.

For dispersing islets, isolated islets were washed with HBSS (without Ca^2+^ or Mg^2+^) and dissociated with 0.5mL of Accutase (innovative cell technology) for 10 minutes at 37°C with intermittent pipetting every 5 minutes. After dissociation, 0.5mL of islet media is added to stop the Accutase activity.

### Hormone secretion assays

5-7 pancreatic islets or dispersed islet cells were seeded into each well of a half-area 96-well plate (Corning Costar) coated with 100μg/mL Rh-Laminin 521 (Gibco) and incubated with islet media overnight before experiments. Islets were washed once with equilibrium media (RPMI 1640, 20mM HEPES, 0.1% BSA, 2mM glucose) and incubated with 50µL of equilibrium media for 45 minutes at 37°C.

After incubation, the equilibration buffer was saved, and islets were incubated with islet secretion buffer (RPMI 1640, 20mM HEPES, 0.1% BSA) with low (1mM) or high (11mM) glucose with the treatment condition for 1 hour at 37°C. After incubation, the secretion buffer was saved. Glucagon and insulin were measured in duplicate with Lumit™ Glucagon and Insulin Immunoassay (Promega), respectively. For normalizing the data with respect to islet mass, the secreted glucagon level was calculated by dividing the glucagon level in the secretion sample by the equilibration value that was divided by the median value of all glucagon levels in equilibration samples. For relative glucagon and insulin secretion level calculation, media for low glucose (1mM, for glucagon secretion) or high (11mM, for insulin secretion) was calculated, and secreted glucagon or insulin level was divided by this median value to get ‘relative secretion’ value.

### Conditioned buffer collection and fractionation

Fully differentiated imBAT cells were briefly washed with PBS and incubated with Krebs-Ringer bicarbonate HEPES (KRBH) buffer (128.8 mM NaCl, 4.8 mM KCl, 1.2 m0M KH_2_PO_4_, 1.2 mM MgSO_4_, 2.5 mM CaCl_2_, 20 mM HEPES, and 5 mM NaHCO_3_, pH 7.4) with 11mM glucose at 37°C for 4 hours and then collected. The collected conditioned buffer was filtered with a 0.22um PES filter (Genesee Scientific) and fractionated with Ultracel 100kDa ultrafiltration Discs (Millipore) using Amicon Stirred Cells system (Millipore) using nitrogen gas pressure (10psi). The fraction with a molecular weight greater than 100kDa (CB-100) was collected and protein concentration was measured using Pierce Quantitative Peptide Assays (Thermo Scientific) for further analysis and experimentation.

For further fractionation, the CB-100 served as the starting material for liquid chromatography experiments. Fractions eluted from an anion-exchange column were collected using an AKTA pure 25M system equipped with a multi-wavelength detector. The column (Cytiva, HiScale 26/20) was packed in-house with Capto Q resin (Cytiva, 17531602). During the binding step, the chromatography method employed buffer A (100 mM Tris, pH 8.0), while the elution step utilized a linear gradient from 0% to 50% of buffer B (100 mM Tris, pH 8.0, 1 M NaCl). To evaluate biological activity, fractions were tested for their ability to inhibit glucagon secretion from pancreatic islets. The active fractions were subsequently concentrated and processed through a pre-packed size-exclusion column (Superdex 200 Increase 10/300 GL), with the running buffer consisting of 100 mM Tris, pH 8.0. Once again, all the fractions were evaluated for their biological activity.

### Size Exclusion Column Peaks Molecular Weight Estimation

To estimate the molecular weight of the elution fractions, protein standard mix (15∼600 kDa; includes thyroglobulin (670 kDa), γ-globulins (150 kDa), albumin (44.3 kDa), and ribonuclease-A (13.7 kDa)) was added to Superdex 200 Increase 10/300 GL column and processed with the same running condition as processing CB-100. The retention factor (K_av_) was calculated using the formula (Elution volume – Void volume)/(Total column volume – Void volume). For the Superdex 200 Increase 10/300 GL column, the total column volume is 23.5 mL, and the reported void volume is 8 mL. A linear regression of log (molecular weight) against K_av_ was used to interpolate the estimated molecular weight of nidogen-2 peaks based on their elution volumes.

### Mass spectrometry sample preparation

Size exclusion chromatography (SEC) fractions were diluted 1:1 with 8 M Urea, 75 mM NaCl, 50 mM Tris, pH 8. The SEC fractions were then reduced with 5 mM dithiothreitol for 45 minutes at 37°C, followed by alkylation for 30 minutes at room temperature with 50 mM chloroacetamide. Then, samples were diluted to 1 M Urea with 50 mM Tris, pH 8, and digested with 0.4 µg of trypsin overnight at 37°C at 800 RPM on a thermomixer. After digestion, samples were quenched with trifluoroacetic acid and dried in a speed-vac. Each SEC fraction was then desalted using C18 ZipTips (Millipore Sigma) prior to LCMS analysis.

### Proteomics data acquisition and analysis

Digested SEC fractions were separated via reverse-phase nano-HPLC using an RSLCnano Ultimate 3000 (Thermo Fisher Scientific). The mobile phase consisted of water + 0.1% formic acid as buffer A and acetonitrile + 0.1% formic acid as buffer B. Peptides were loaded onto a µPAC□□ Trapping column (PharmaFluidics) and separated on a 50 cm µPAC□□ column (PharmaFluidics) operated at room temperature and flowing at 750 or 300 nL/min using a segmented gradient as follows: 5 min from 2-12% buffer B at 750nl/min, then 42 min from 12-20% B at 300nl/min, and finally 16 min from 20-40% B at 300nl/min. Mass spectrometry analysis was performed on an Orbitrap Eclipse (Thermo Fisher Scientific) using a data-independent acquisition method. MS1 scans were obtained at 60,000 resolution in the Orbitrap with a scan range from 390-1010 m/z, 60 ms max injection time, and an automated gain control (AGC) target of 1e6. Cycles of 38 DIA MS2 scans were obtained at 30,000 resolution in the Orbitrap, using 16 m/z wide quadrupole isolation windows, 54ms max injection time, 5e5 AGC target, and 30% higher-energy collision dissociation energy.

After data acquisition, peptide and protein identification and quantification were performed by DIA-NN (1.8.2 beta 27). An in silico tryptic spectral library was created for the mouse SwissProt proteome (downloaded 8/27/19), along with common contaminants for a total of 17,194 protein entries. Carbiomothemylated cysteine was used as a static modification, one oxidized methionine was set as a variable modification, and one missed cleavage was allowed. Heuristic protein inference was enabled, all protein and gene identifications were globally controlled to an FDR of 0.01, quantification was performed with QuantUMS (high precision), and normalization was disabled. Secreted protein annotations were downloaded from The Human Protein Atlas^56^. The mass spectrometry proteomics data have been deposited to the ProteomeXchange Consortium via the PRIDE partner repository.

### Extracellular vesicles (EVs) isolation and particle counting

Sepharose CL-2B particles (Sigma-Aldrich, CL2B300) were packed a few hours before use. CL-2B slurry was prepared by mixing 75% beads with 25 % of 50% aqueous isopropanol. A 10mL centrifuge column (Pierce, 89898) was prepared by rinsing the column with 50% isopropanol and slowly adding 15mL of CL-2B slurry to settle down and condense as the filter retains the beads. Once beads are settled, the column was washed with 13mL of 50% isopropanol twice and DI water once. CL-2B size exclusion column was conditioned by running 50mL of PBS through the column. For separation of the EVs fraction, 1mL of CB-100 was added to the conditioned CL-2B column, and elution was fractionated (0.5mL) by adding 1.5 column volume of PBS. Each fraction was tested for glucagon secretion inhibition activity, counted for EVs particle number with NTA ZetaView, and measured for absorbance at 205nm with a Nanodrop.

### Live-cell imaging

Pancreatic islets were seeded onto an 8-well-chambered cover glass system (Cellvis) coated with Rh-laminin521 (Gibco, 2μg/cm^2^) and incubated in islet media for 3 days prior to experiments. For Ca^2+^ imaging, the seeded islets were placed in a Zeiss LSM 880 confocal microscope with a heated, CO2-controlled microscope stage (37°C, 5% CO2) and a Plan-Apochromat 63X 1.4 NA oil immersion objective. The islet media was switched to imaging media (KRBH with 0.1% BSA) with 2 mM glucose for 30 minutes. For secretion, the buffer was changed to islet media with or without insulin receptor antagonist (S961) (Phoenix Pharmaceuticals) in 1mM or 11mM glucose and incubated for 20 minutes at 37°C, 5% CO2. After incubation, an initial baseline time course image was acquired for 15 minutes. After baseline image acquisition, the buffer was switched to imaging media with either 1mM or 11mM glucose with treatment (CB-100: 5μg/mL, nidogen-2 (40ng/mL, Biotechne R&D Systems)) and incubated for 5 minutes. After 5 minutes, the treatment image time course was acquired for 15 minutes. GCaMP6f fluorescence was acquired with 488nm laser excitation, detected at an emission range of 490-597nm with the spectral detector as described ^54^.

For cAMP imaging, Epac^SH187^–cAMP FRET biosensor (mTurquoise2Δ_Epac(CD,Δ DEP,Q270E)_cpVenus_Venus) was used ^57^. The sensor was packaged in adenovirus for delivery, and virus particles were concentrated to a titer of 3.28×10^12^ particles/mL. Adenovirus stock is directly added to islet culture media (3.28×10^11^ particles/mL) and cultured for 3 days at 37°C, 5% CO_2_ prior to the experiment. The seeded islets were placed in a Zeiss LSM 880 confocal microscope with a heated, CO2-controlled microscope stage (37°C, 5% CO2) and a Plan-Apochromat 20X/0.2 M27 objective. Islets were equilibrated for 30 minutes in imaging media with 2mM Glucose. The baseline image was acquired after incubating islets with KRBH with 0.1% BSA with or without S961 (Phoenix Pharmaceuticals Inc) in 1 or 11mM glucose for 30 minutes. Then, the buffer was switched to imaging media with either 1mM or 11mM glucose with treatment (CB-100: 5μg/mL, nidogen-2 (Biotechne R&D Systems)) and incubated for 30 minutes prior to acquiring a treatment image. RFP α-cells were imaged with 561nm laser excitation to mark the RFP α-cells. FRET signals for acceptors and donors (mVenus, mTurquoise) were measured with LSM 880 spectral detector mode with 405 nm excitation and detected at an emission range of 445-650 nm. Using the entire emission spectrum image, the acceptor and donor’s signals were acquired through linear unmixing with the ZEISS ZEN imaging software. For analysis, the percentage change of the Donor-to-acceptor ratio was calculated before and after treatment (the FRET signal is inversely proportional to the concentration of cAMP).

### CB-100 and nidogen-2 injection *in vivo* experiment

NOD mice with blood glucose levels between 190mg/dL and 230mg/dL received daily injections of CB-100 (1.5 mg/kg BW, nape of the neck, subcutaneous) or sham (KRBH) for 7 consecutive days on the same time. Blood glucose levels were measured weekly with a blood glucose meter (Glucocard Vital). After 2∼8 weeks of blood glucose monitoring (measurement time around 13:00), mice underwent IPGTT, IPITT, and metabolic activity studies, and were euthanized for tissue sample collection.

### Glucose and insulin tolerance tests

Mice underwent a 6-hour fast before IPGTT or a 4-hour fast (shorter fasting time to prevent severe hypoglycemia from insulin injection) before IPITT. After a brief isoflurane anesthesia, blood glucose levels and body weight were recorded, and mice received intraperitoneal injections of 0.75 IU/kg BW Humulin R (Lilly) (for IPITT) or 2g/kg BW glucose (for IPGTT). After the injection, blood glucose levels were measured at 15, 30, 60, 90, and 120 minutes.

### Body composition analysis

After measuring the body weight, lean and fat mass was measured with Echo MRI (EchoMRI) without anesthesia as previously described ^58^. Two measurements were averaged to calculate the % mass with the body weight.

### Metabolic studies

Mice were housed in the TSE Phenomaster system (TSE Systems) overnight before the data was collected. After overnight acclimation, metabolic activities (heat, VO_2_, and activity) data was collected as previously outlined ^59^. Metabolic activities were measured every 15 minutes for 3 days, with four measurements averaged for each hourly data point.

### Tissue sample collection from *in vivo* study

Following weekly blood glucose measurements around 13:00, animals were euthanized, and blood plasma was collected via cardiopuncture. The collected blood was mixed with 15 μL of EDTA and centrifuged at 2000g for 15 minutes at 4°C. The supernatant (blood plasma) was then collected and stored at −80°C for further analysis. The entire pancreas, inguinal WAT, BAT, and gastrocnemius muscle were collected, while the liver underwent perfusion with PBS before collection.

### Hormone analysis

Glucagon and insulin were measured with ELISA kit (Crystal Chem) for blood plasma hormone measurement. For Pancreatic hormone analysis, pancreatic hormones were extracted to 2mL of acid ethanol per mouse following previously reported methods ^15^. Pancreatic insulin and glucagon levels were measured with Immunoassay kits (Promega). If the entire pancreas was used to extract pancreatic hormone levels, the mice body weight was used for normalization. If a portion of the pancreas was used, tissue weight or protein concentration was used for normalization.

### Histology

For histological analysis, the collected tissues were briefly washed with cold PBS, fixed in 10% Neutral buffered formalin for 2 days at room temperature, and transferred to 70% ethanol for at least 2 days. For hematoxylin-eosin (H&E) staining and immunofluorescence (IF) staining, neutral buffered formaldehyde-fixed tissues were submitted for sectioning to the Anatomic and Molecular Pathology (AMP) Core Labs at Washington University in St. Louis for the paraffin embedding and sectioning into 5um cross-sections. H&E staining was also performed by the AMP core. The paraffin embedded sections were deparaffined (xylene, 10min, 5min), rehydrated (100% EtOH (10min), 90% EtOH (5min), 70% EtOH (5min), PBS (5min)), boiled with antigen retrieval (Diva Decloaker, 15min), permeabilized with PBS-T (PBS with 0.1% Tween 20; 1hour), and blocked (PBS with 0.05% Tween 20 and 2% BSA, 1hour). For adhered cell IF imaging, cells were fixed with 4% PBS buffered formalin for 30 minutes and permeabilized with PBS with 0.1% Triton X-100 for 15 minutes. Then cells were blocked with PBS with 0.05% Tween-20 and 2% BSA for 30 minutes.

For immunofluorescence staining, the sample was incubated with primary antibodies diluted in a blocking buffer at 4°C overnight (See Supplementary Table S2 for detailed antibody information). After washing slides with PBS 3 times (10min, 5min, 5min), slides were incubated with secondary antibodies in room temperature for 1.5 hours, DAPI staining for 5 minutes, washed, and mounted with prolong glass antifade Mountant (Invitrogen). The image was acquired with Carl Zeiss LSM 880 Airyscan two-photon confocal microscope.

### RNA sequencing and analysis

RNA from the tissue or cells was isolated using RNeasy Lipid mini (Qiagen). Isolated RNA samples were submitted to The Genome Access Technology Center at Washington University in St. Louis for the RNA sequencing. All gene counts were imported into the R DESeq2 package. For cluster analysis, ComplexHeatmap package was used (top 15 most upregulated or downregulated genes were represented; cutoff: p_adjust < 0.05, abs (Log_2_FC) > 2, basemean > 50). For enrichment analysis, clusterProfiler package using biomaRt and pathfindR package (Log_2_FC > 2 cutoff) were used, and top 10 more enriched processes were represented. For gene-set based analysis, DEGs entrezgene from BAT (CB-100 treated vs sham; Log_2_FC >2) was entered to ConsensusPathDB-mouse (http://cpdb.molgen.mpg.de/MCPDB) ^60^ (Reactome and kEGG pathway, minimum overlap with input list :2, p-value cutoff :0.01). For gene ontology categories, level 2 and level 3 were selected for all biological, molecular, and cellular components with p-value cutoff of 0.01. Top 5 reactome and KEGG-enriched pathways are represented. To generate a volcano plot, the EnhancedVolcano package was used. For ingWAT, p_adjust < 0.0005, abs (Log_2_FC) > 5 cut-off was used. For BAT p_adjust < 0.05, abs (Log_2_FC) > 3 cutoff was used.

### Quantitative PCR analysis

RNA was extracted from tissues or cells using either the RNeasy lipid mini kit (Qiagen) or TRIzol reagent (Invitrogen), and cDNA was synthesized using the Superscript VILO cDNA synthesis kit (Invitrogen). Quantitative real-time polymerase chain reaction (PCR) was conducted using the PowerUp SYBR Green master mix (Applied Biosystems) and the ViiA7 Real-Time PCR System (Applied Biosystems). The comparative cycle-threshold (Ct) method was used to calculate relative mRNA expression level; ΔCt value was calculated with housekeeping gene (18S) and euglycemic mice or untreated cell’s ΔCt value were used as a control sample or euglycemic mouse for ΔΔCt calculation. Primers for cDNA amplification are detailed in Supplementary Table S3.

### *In vitro* glucose consumption assay

Pre-existing culture media were removed, and the cells were briefly washed once with PBS before introducing fresh culture media (DMEM with 10% FBS and 1% penicillin-streptomycin) and incubated overnight. After overnight incubation, 10µL of culture media was collected (pre-testing culture media). Cells were briefly washed once with PBS before the introduction of fresh culture media (DMEM with 10% FBS and 1% penicillin-streptomycin) with or without the following treatments: CB-100 (52.8 µg/mL), nidogen-2 (5 µg/mL), insulin (10 nM), and S961 (200 nM). The cells were then incubated for 24 hours at 37°C and 5% CO_2_. After incubation, the culture media were collected.

The glucose concentration in the pre-testing, pre-incubation, and post-incubation culture media was measured using a high-sensitivity glucose assay (Abcam). To account for differences in cell numbers across wells, the glucose consumed in the pre-testing culture media in each well was normalized by dividing by the median consumed glucose level in all wells. This value represented the glucose consumption-ability of each well. Consumed glucose was calculated by (pre-incubation media – post-incubation media) / glucose-consumption ability for each well. To determine the relative glucose consumption, the consumed glucose for each condition was divided by the median consumed value of the control (untreated) condition.

### Western blotting

CB-100 was mixed with 4X protein sample loading buffer (Li-COR) and boiled for 5 minutes at 95°C. Prepared samples were used for electrophoresis with 4-20% protein gel (Bio-Rad) and transferred to a PVDF membrane (Millipore). After blocking for 1 hour with TBS-T (TBS with 0.1% Tween20) with 2% BSA, the membrane was incubated with Nidogen-2 polyclonal Rabbit (1:1000, ProteinTech) in TBS-T with 1% BSA at 4°C overnight with shaking. Then, the membrane was washed with TBS-T 3 times and incubated with a IRDye® 680RD Goat anti-Rabbit IgG Secondary Antibody (1:2000, LI-COR) in TBS-T with 1% BSA for an hour at room temperature with shaking. The membrane was washed with TBS-T 3 times (10 minutes each) and visualized by Li-COR Odyssey M Imager (Li-COR).

Protein lysates were extracted from L6-GLUT4myc myotubes with 1% Nonidet P-40 lysis buffer containing 25 mM HEPES (pH 7.4), 1% Nonidet P-40, 10% glycerol, 50 mM sodium fluoride, 10 mM sodium pyrophosphate, 137 mM NaCl, 1 mM sodium vanadate, 1 mM phenylmethylsulfonyl fluoride, 10 μg/ml aprotinin, 1 μg/ml pepstatin, and 5 μg/ml leupeptin. Boiled protein lysates with 1x Laemmli SDS sample buffer (Bioland Scientific) were resolved using 8% sodium dodecyl sulfate-polyacrylamide gels (SDS-PAGE) with pre-stained molecular weight markers. Subsequently, the resolved protein lysates were transferred to nitrocellulose or polyvinylidene difluoride membranes (PVDF) and were stained with Ponceau S. The membranes were then sectioned into small strips based on the molecular weight of each protein for immunoblotting. Immunoreactive bands were detected using enhanced chemiluminescence (ECL) or ECL Prime reagents (Cytiva Amersham™ ECL™). The detection process was followed by imaging the immunoreactive bands using a Chemi-Doc Touch gel documentation system (Bio-Rad). The primary antibodies used in this study include AKT (1:2000, Cell Signaling), p-AKT (1:2000, Cell Signaling), IRS-1 (1:1000, Cell Signaling), and p-IRS-1 (1:1000, Millipore Sigma). For the secondary antibody (dilution-1:10,000), Goat Anti-Rabbit IgG (H L)-HRP Conjugate or Goat Anti-Mouse IgG (H L)-HRP Conjugate (Bio-Rad) was incubated with the blot for 1□h at room temperature (RT). The quantification of the signals was carried out using Image Lab v. 5.1 software and normalized to Ponceau. Original western blot images have been submitted as the supplementary figure.

## STATISTICAL ANALYSIS

Data were analyzed with R studio, Microsoft Excel, Image J, MATLAB, and GraphPad Prism. Calcium images were analyzed with MATLAB as described ^54^. The statistical significance of each data is reported as mean ± SEM. Statistical significance was calculated as described in each figure legend.

## SUPPLEMENTAL FIGURE TITLES AND LEGENDS

**Supplementary Figure S1. Further Characterization of Large Brown Adipocyte-Secreted Proteins (CB-100). Related to Figure 1 and Figure 6**.

(**A**) Relative insulin secretion measured in low (yellow, 1mM) and high (teal, 11mM) glucose conditions in both human (non-diabetic n=16, T1D n=1, T2D n=7, see Supplementary Table S1 for more detail) and mouse (non-diabetic n=30, T1D model n=5, T2D model n=3) pancreatic islets.

(**B**) Relative mRNA expression of *Cidea* and *Ucp1* of imBAT without (pink) or with (green) norepinephrine stimulation (10μM, 16 hours).

(**C**) Relative glucagon secretion in low (yellow, 1mM) and high (teal, 11mM) glucose conditions in pancreatic islets from non-diabetic mice (n=4) with CB-100 (26.5μg/mL) collected from imBAT without (NE-) or with (NE+) stimulation (10μM, 16 hours).

(**D**) Relative glucagon secretion measured without (navy blue) or with CB-100 (purple, 5μg/mL) in low (1mM) glucose condition with GLP-1 receptor (GLP-1R) antagonist (Exendin-3, 50nM) or glucagon receptor (GCGR) antagonist (Adomeglivant, 1.5μM) or Somatostatin receptor (SSTR) antagonist (CYN154806, 25nM) or Ephrin A4 receptor (EphA4R) antagonist (KYL, 50μM), insulin receptor (INSR) antagonist (S961, 1μM) in non-diabetic mouse pancreatic islets (n=3). Related to Figure 6B.

Data are presented as Mean ± SEM. Statistical significance was determined using an unpaired t-test.

Groups compared for statistical analysis are indicated by the line. *p < 0.05, **p < 0.01, ***p < 0.001, ns (p > 0.05).

**Supplementary Figure S2. Detailed Characterization of CB-100 Treated NOD Mice. Related to Figure 2 and Figure 4**

(**A&D**) CB-100 treatment success (green) and failed (red) percentage of early (female: n=21, male: n=12, blood glucose (BG) between 190mg/dL and 230mg/dL) and late-stage (female: n=14, male: n=5, >230mg/dL) of hyperglycemia in **(A)** female and **(D)** Male NOD mice. Related to Figure 2B.

(**B**) Weekly non-fasting blood glucose levels following seven days of injection of CB-100 (subcutaneous, 1.5 mg/kg BW; blue, success n=19 & reverted n=2) and sham (KRBH, black, n = 12) female NOD mice from late hyperglycemia (18-22 weeks old, BG between 190mg/dL and 230mg/dL). Related to Figure 2B.

(**C**) Weekly non-fasting blood glucose levels following seven days of injection of CB-100 (subcutaneous, 1.5 mg/kg BW; Success, green, n = 3 & failed, red, n=11) and sham (KRBH, black, n = 6) female NOD mice from late hyperglycemia (18-22 weeks old, BG >230mg/dL). Related to Figure 2B.

**(E)** Weekly non-fasting blood glucose levels following seven days of injection of CB-100 (subcutaneous, 1.5 mg/kg BW; blue, success n=11 & reverted n=1) and sham (KRBH, black, n = 6) male NOD mice from late hyperglycemia (18-22 weeks old, BG between 190mg/dL and 230mg/dL). Related to Figure 2B.

**(F)** Weekly non-fasting blood glucose levels following seven days of injection of CB-100 (subcutaneous, 1.5 mg/kg BW; green, success n=1 & failed, red, n=4) and sham (KRBH, black, n = 12) male NOD mice from late hyperglycemia (18-22 weeks old, BG >230mg/dL). Related to Figure 2B.

**(G-I)** Metabolic cages measured and averaged (**F**) VO_2_ consumption, (**G**) activity, and (**H**) heat (kcal/h/kg) from euglycemic control (yellow, n = 2), sham (red, n = 12), and CB-100-treated (brown, n = 4) NOD mice. Related to Figure 2C.

(**J**) H&E Staining of liver sections from euglycemic control, sham-treated, and CB-100 treated NOD mice Scale bar: 100 μm. Related to Figure 4

(**K**) Immunofluorescence of GLUT4 (green) and nuclei (blue) in brown adipose tissue of euglycemic control, sham-treated, and CB-100-treated NOD mice. Scale bar: 100 μm. Related to Figure 4

(**L**) Relative expression levels of trafficking Regulator of GLUT4 1 (*Trarg1*) and glucose transporter type 4 (*Glut4*) in BAT of euglycemic control (green, n=5) sham-treated (red, n=5) and CB-100-treated (yellow, n=7) NOD mice. Related to Figure 4.

Data are presented as Mean ± SEM. Statistical significance was determined using an unpaired t-test. Groups compared for statistical analysis are indicated by the line. *p < 0.05, **p < 0.01, ***p < 0.001, ns (p > 0.05).

**Supplementary Figure S3. CB-100 *in vitro* glucose uptake and western blotting analysis. Related to Figure 4**.

(**A**) Immunoblot (IB) for p-IRS-1(Tyr^612^), total IRS-1, and p-AKT(Ser^463^) and total AKT normalized with Ponceau staining in fully differentiated L6-GLUT4myc myotube cells untreated (Vehicle) and treated with CB-100 (10μg/mL, 16 hours) and spiked in insulin (10nM, 5minutes). Related to Figure 5G.

(**B**) Relative glucose consumption in myotubes (C2C12), hepatocytes (AML12), white adipocytes (3T3-L1), and brown adipocytes (imBAT) treated without (control, navy blue) or with insulin (red, 10nM) and in the absence (control) or presence of S961 (200 nM) for 24 hours.

(**C**) IB for p-IRS-1(Tyr^612^), total IRS-1, and p-AKT(Ser^463^) and total AKT normalized with Ponceau staining in fully differentiated L6-GLUT4myc myotube cells untreated (Vehicle) and treated with insulin (10nM, 16∼18 hours), spiked in insulin (10nM, 5 minutes).

(**D**) IB analysis of p-IRS-1^Tyr606^/total-IRS-1 and p-AKT^Ser^/total-AKT from L6-GLUT4myc myotube untreated (navy blue, control) or treated with insulin (red, 10nM, 16∼18 hours) then spiked in insulin (10nM, 5minutes).

Data are presented as Mean ± SEM. Statistical significance was determined using an unpaired t-test. Groups compared for statistical analysis are indicated by the line. *p < 0.05, **p < 0.01, ***p < 0.001, ns (p > 0.05).

**Supplementary Figure S4. Candidate Protein Testing and Further Investigation of the Effects of Nidogen-2 on Islets Secretion and *in vitro* Glucose uptake. Related to Figure 6**.

**(A, C & E)** Relative glucagon secretion was measured in low (yellow, 1mM) and high (teal, 11mM) glucose conditions in non-diabetic mice (n=6) without or with (**A)** nidogen-2 (Nid2) (**C)** superoxide dismutase 3 (Sod3) or (**E**) complement factor H (CFH). Concentration with a significant inhibition activity was highlighted with red.

**(B&D)** Glucagon secretion level with SEC fractions at low glucose (1mM) condition (grey bar) plotting with protein intensity (red) of (**B**) superoxide dismutase 3 (Sod3) and (**D**) complement factor H (CFH). Glucagon secretion of a control (1mM and 11mM) is indicated with the left two dark grey bars. The expected lowest glucagon secretion was determined with the glucagon secretion level at high glucose condition (11mM) and marked with the dotted line.

(**F**) Relative glucose consumption measured in C2C12, AML12, 3T3-L1, and imBAT cell line without (control, navy blue) or with nidogen-2 (orange, 1μg/mL) (16 hours).

(**G**) Relative glucose consumption in imBAT treated without (control, navy blue) or with nidogen-2 (orange, 1μg/mL) and in the absence (control) or presence of S961 (200 nM) for 24 hours.

Data are presented as Mean ± SEM. Statistical significance was determined using an unpaired t-test. Groups compared for statistical analysis are indicated by the line. *p < 0.05, **p < 0.01, ***p < 0.001, ns (p > 0.05).

