## Supplementary Figure and tables for "Insulin-Independent Regulation of Type 1 Diabetes via Brown Adipocyte-Secreted Proteins and the Novel Glucagon Regulator Nidogen-2"

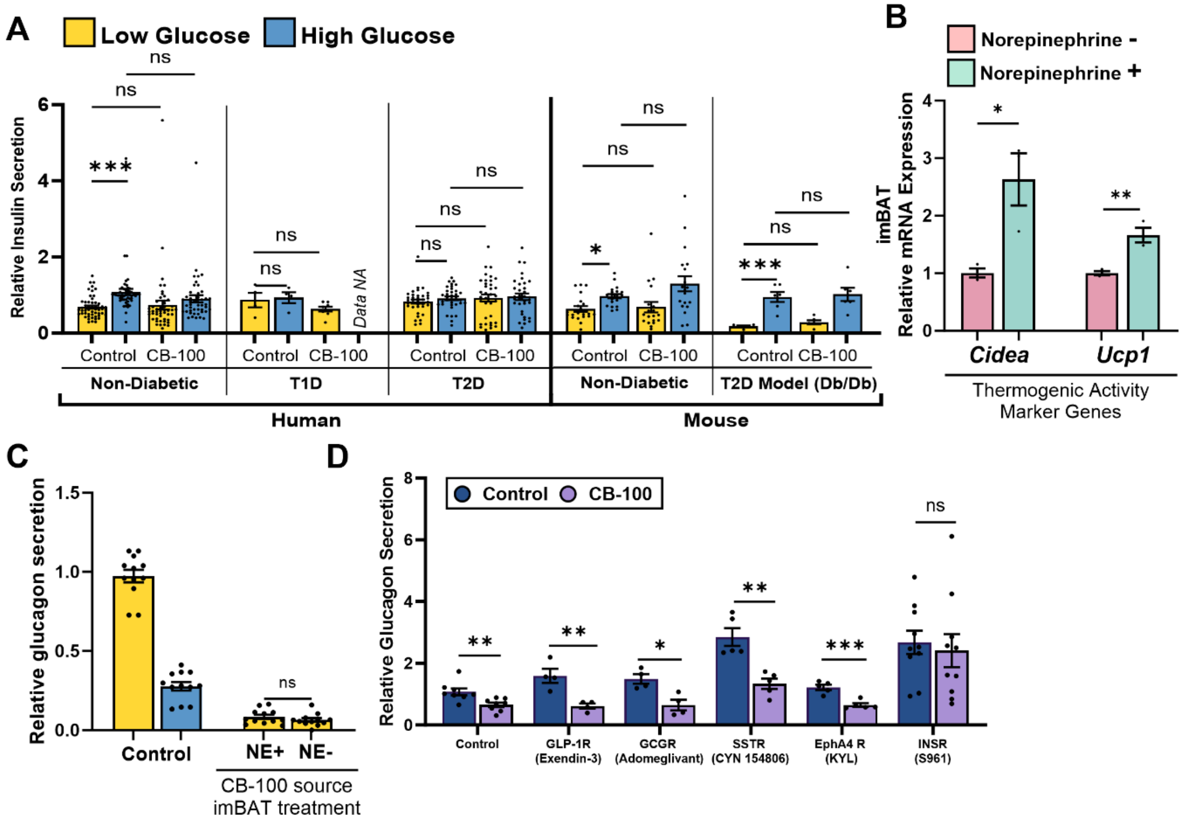


**Supplementary Figure S1. Further Characterization of Large Brown Adipocyte-Secreted Proteins (CB-100). Related to Figure 1 and Figure 6.**

(**A**) Relative insulin secretion measured in low (yellow, 1mM) and high (teal, 11mM) glucose conditions in both human (non-diabetic n=16, T1D n=1, T2D n=7, see Supplementary Table S1 for more detail) and mouse (non-diabetic n=30, T1D model n=5, T2D model n=3) pancreatic islets.

(**B**) Relative mRNA expression of *Cidea* and *Ucp1* of imBAT without (pink) or with (green) norepinephrine stimulation (10µM, 16 hours).

(**C**) Relative glucagon secretion in low (yellow, 1mM) and high (teal, 11mM) glucose conditions in pancreatic islets from non-diabetic mice (n=4) with CB-100 (26.5µg/mL) collected from imBAT without (NE-) or with (NE+) stimulation (10µM, 16 hours).

(**D**) Relative glucagon secretion measured without (navy blue) or with CB-100 (purple, 5µg/mL) in low (1mM) glucose condition with GLP-1 receptor (GLP-1R) antagonist (Exendin-3, 50nM) or glucagon receptor (GCGR) antagonist (Adomeglivant, 1.5µM) or Somatostatin receptor (SSTR) antagonist (CYN154806, 25nM) or Ephrin A4 receptor (EphA4R) antagonist (KYL, 50µM), insulin receptor (INSR) antagonist (S961, 1µM) in non-diabetic mouse pancreatic islets (n=3). Related to Figure 6B.

Data are presented as Mean ± SEM. Statistical significance was determined using an unpaired t-test. Groups compared for statistical analysis are indicated by the line. *p < 0.05, **p < 0.01, ***p < 0.001, ns (p > 0.05).

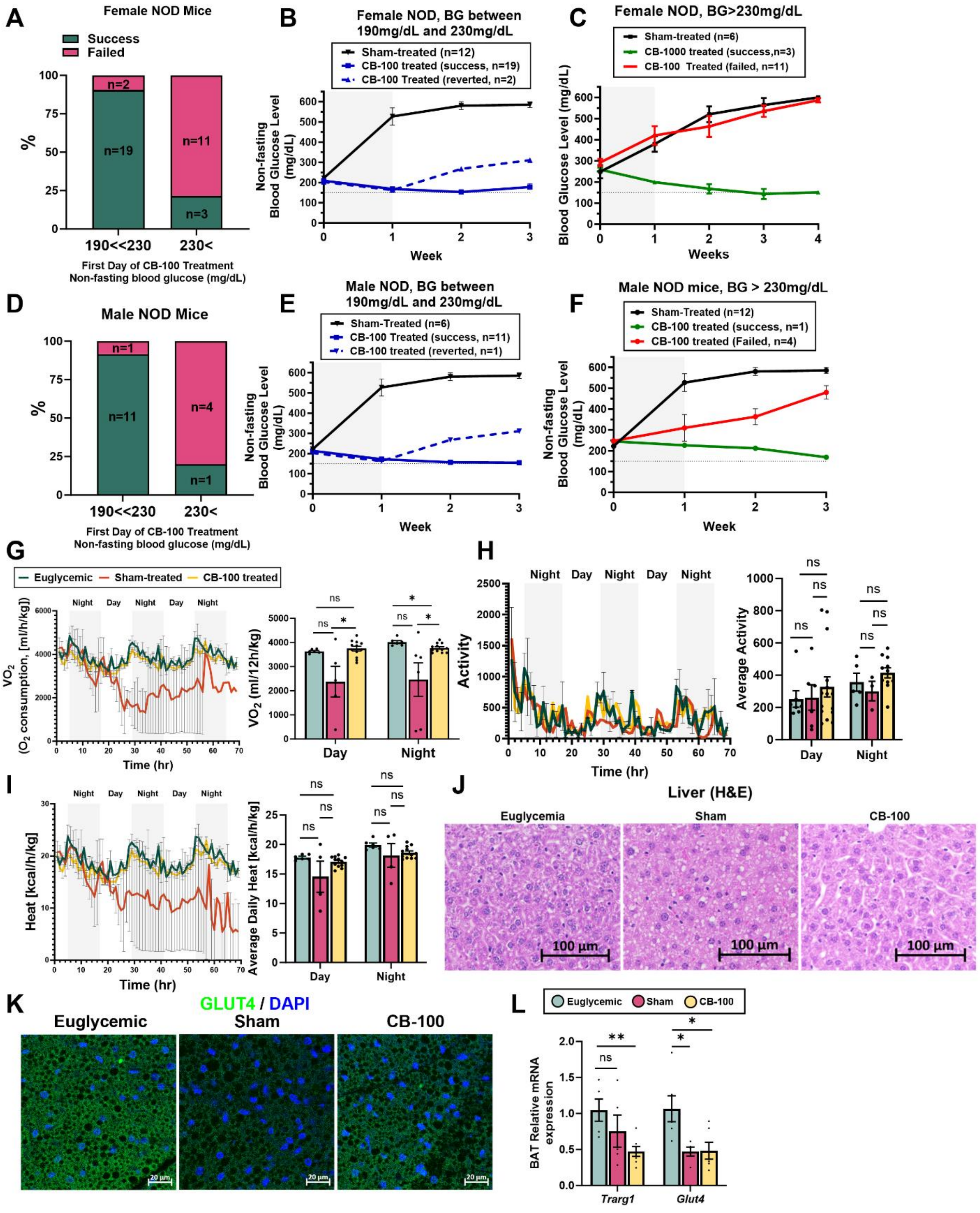

A
Female NOD Mice
Success
Failed
n=2
n=19
n=11
n=3
%
190<<230
230<
First Day of CB-100 Treatment
Non-fasting blood glucose (mg/dL)
B
Female NOD, BG between 190mg/dL and 230mg/dL
Sham-treated (n=12)
CB-100 treated (success, n=19)
CB-100 Treated (reverted, n=2)
Non-fasting Blood Glucose Level (mg/dL)
Week
C
Female NOD, BG>230mg/dL
Sham-treated (n=6)
CB-1000 treated (success,n=3)
CB-100 Treated (failed, n=11)
Blood Glucose Level (mg/dL)
Weeks
D
Male NOD Mice
n=1
n=11
n=4
n=1
%
190<<230
230<
First Day of CB-100 Treatment
Non-fasting blood glucose (mg/dL)
E
Male NOD, BG between 190mg/dL and 230mg/dL
Sham-Treated (n=6)
CB-100 Treated (success, n=11)
CB-100 treated (reverted, n=1)
Non-fasting Blood Glucose Level (mg/dL)
Week
F
Male NOD mice, BG > 230mg/dL
Sham-Treated (n=12)
CB-100 treated (success, n=1)
CB-100 treated (Failed, n=4)
Non-fasting Blood Glucose Level (mg/dL)
Week
G
Euglycemic
Sham-treated
CB-100 treated
Night
Day
VO2 (O2 consumption, [ml/h/kg])
Time (hr)
VO2 (ml/12h/kg)
Day
Night
H
Activity
Time (hr)
Average Activity
I
Heat [kcal/h/kg]
Time (hr)
Average Daily Heat [kcal/h/kg]
J
Liver (H&E)
Euglycemia
Sham
CB-100
100 µm
K
GLUT4 / DAPI
Euglycemic
Sham
CB-100
20 µm
L
Euglycemic
Sham
CB-100
BAT Relative mRNA expression
Trarg1
Glut4

Data are presented as Mean ± SEM. Statistical significance was determined using an unpaired t-test. Groups compared for statistical analysis are indicated by the line. *p < 0.05, **p < 0.01, ***p < 0.001, ns (p > 0.05).

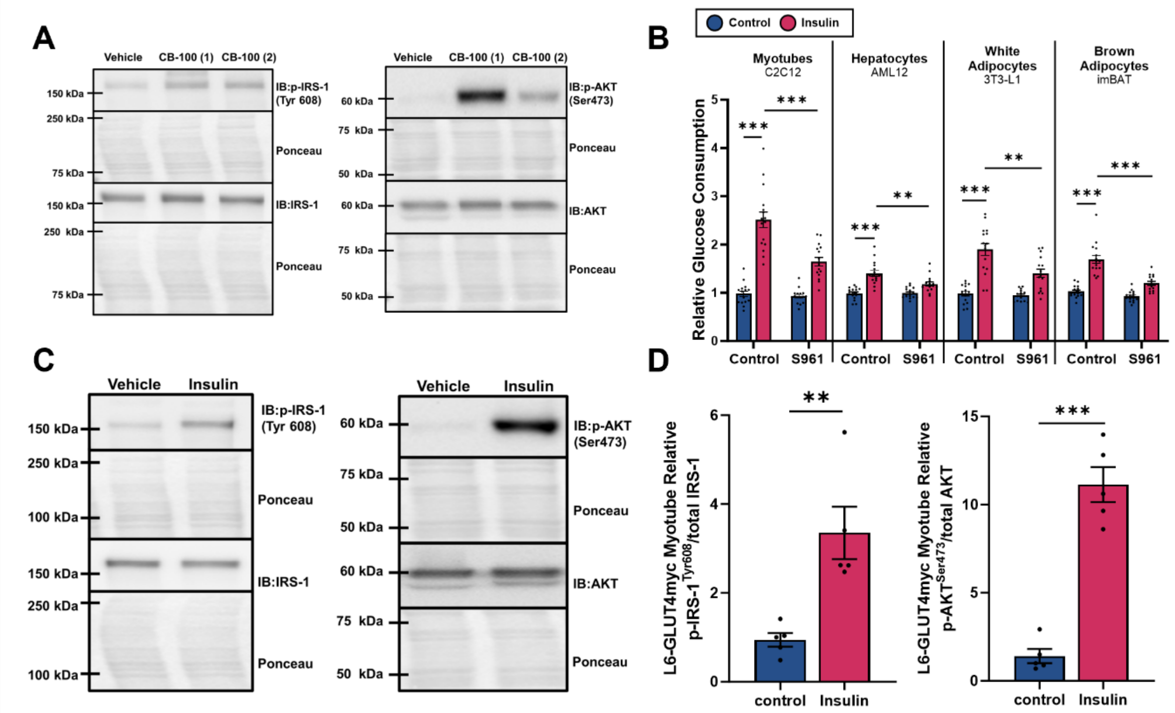


**Supplementary Figure S3. CB-100 *in vitro* glucose uptake and western blotting analysis. Related to Figure 4.**

**(A)** Immunoblot (IB) for p-IRS-1(Tyr$^{612}$), total IRS-1, and p-AKT(Ser$^{463}$) and total AKT normalized with Ponceau staining in fully differentiated L6-GLUT4myc myotube cells untreated (Vehicle) and treated with CB-100 (10µg/mL, 16 hours) and spiked in insulin (10nM, 5minutes). Related to Figure 5G.

(**B**) Relative glucose consumption in myotubes (C2C12), hepatocytes (AML12), white adipocytes (3T3-L1), and brown adipocytes (imBAT) treated without (control, navy blue) or with insulin (red, 10nM) and in the absence (control) or presence of S961 (200 nM) for 24 hours.

**(C)** IB for p-IRS-1(Tyr$^{612}$), total IRS-1, and p-AKT(Ser$^{463}$) and total AKT normalized with Ponceau staining in fully differentiated L6-GLUT4myc myotube cells untreated (Vehicle) and treated with insulin (10nM, 16~18 hours), spiked in insulin (10nM, 5 minutes).

**(D)** IB analysis of p-IRS-1$^{Tyr606}$/total-IRS-1 and p-AKT$^{Ser473}$/total-AKT from L6-GLUT4myc myotube untreated (navy blue, control) or treated with insulin (red, 10nM, 16~18 hours) then spiked in insulin (10nM, 5minutes).

Data are presented as Mean ± SEM. Statistical significance was determined using an unpaired t-test. Groups compared for statistical analysis are indicated by the line. *p < 0.05, **p < 0.01, ***p < 0.001, ns (p > 0.05).

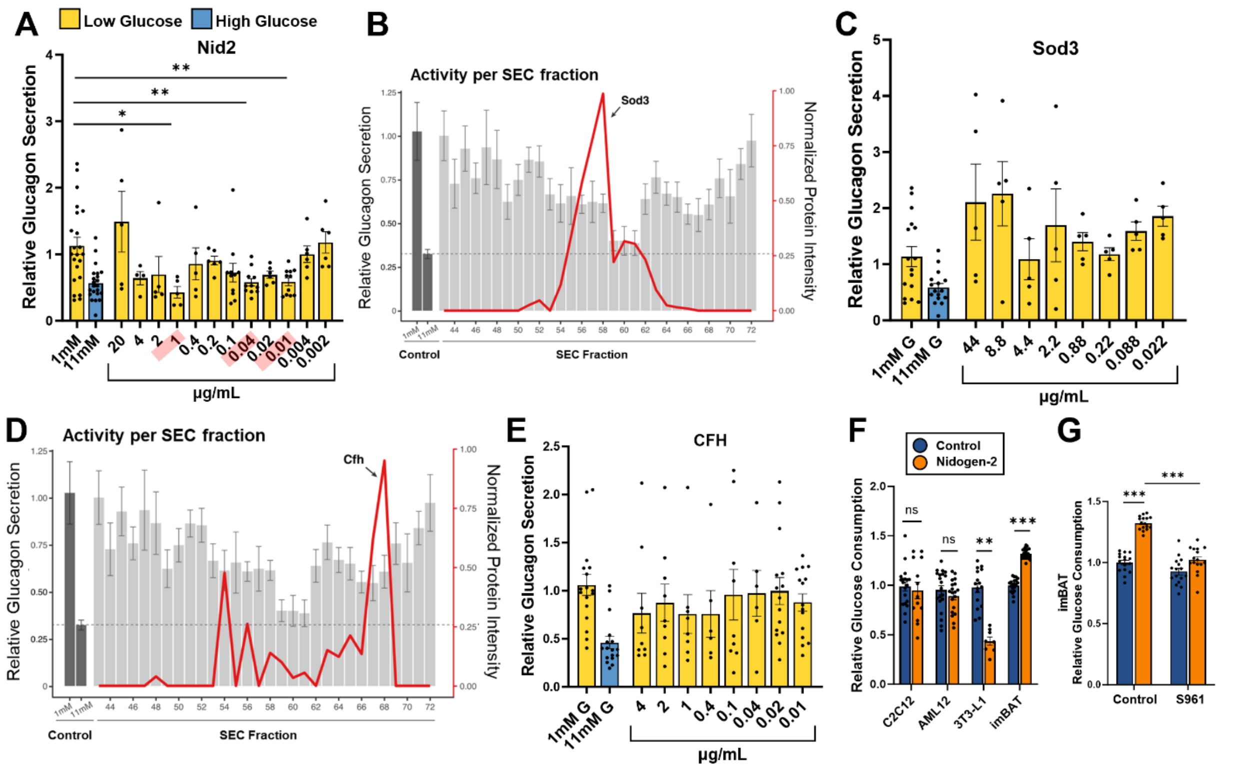


**Supplementary Figure S4. Candidate Protein Testing and Further Investigation of the Effects of Nidogen-2 on Islets Secretion and *in vitro* Glucose uptake. Related to Figure 6.**

**(A, C & E)** Relative glucagon secretion was measured in low (yellow, 1mM) and high (teal, 11mM) glucose conditions in non-diabetic mice (n=6) without or with (**A**) nidogen-2 (Nid2) (**C**) superoxide dismutase 3 (Sod3) or (**E**) complement factor H (CFH). Concentration with a significant inhibition activity was highlighted with red.

**(B&D)** Glucagon secretion level with SEC fractions at low glucose (1mM) condition (grey bar) plotting with protein intensity (red) of (**B**) superoxide dismutase 3 (Sod3) and (**D**) complement factor H (CFH). Glucagon secretion of a control (1mM and 11mM) is indicated with the left two dark grey bars. The expected lowest glucagon secretion was determined with the glucagon secretion level at high glucose condition (11mM) and marked with the dotted line.

(**F**) Relative glucose consumption measured in C2C12, AML12, 3T3-L1, and imBAT cell line without (control, navy blue) or with nidogen-2 (orange, 1μg/mL) (16 hours).

**(G)** Relative glucose consumption in imBAT treated without (control, navy blue) or with nidogen-2 (orange, 1μg/mL) and in the absence (control) or presence of S961 (200 nM) for 24 hours.

Data are presented as Mean ± SEM. Statistical significance was determined using an unpaired t-test. Groups compared for statistical analysis are indicated by the line. *p < 0.05, **p < 0.01, ***p < 0.001, ns (p > 0.05).

**Supplementary Table S1. List of Human Islets Donor Information. Related to Figures 1C, 1D, 5H, 5I, 6A, and S1A.**

| | Donor (RRID) | Age (years) | Sex | BMI | Disease |
|---|---|---|---|---|---|
| 1 | SAMN25519947 | 32 | Male | 24.3 | Non-diabetic |
| 2 | SAMN26646319 | 20 | Male | 20.5 | Non-diabetic |
| 3 | SAMN27361473 | 34 | Male | 31.8 | Non-diabetic |
| 4 | SAMN29771427 | 44 | Male | 27.9 | Non-diabetic |
| 5 | SAMN32641506 | 16 | Male | 29.5 | Non-diabetic |
| 6 | SAMN38518088 | 57 | Female | 30.1 | Non-diabetic |
| 7 | SAMN38334492 | 51 | Male | 24.6 | Non-diabetic |
| 8 | SAMN38226661 | 37 | Female | 30 | Non-diabetic |
| 9 | SAMN37973608 | 55 | Female | 31.7 | Non-diabetic |
| 10 | SAMN37871873 | 59 | Male | 41.7 | Non-diabetic |
| 11 | SAMN36823227 | 41 | Female | 38.4 | Non-diabetic |
| 12 | SAMN35301006 | 52 | Male | 31.6 | Non-diabetic |
| 13 | SAMN34411471 | 45 | Male | 33.1 | Non-diabetic |
| 14 | SAMN34033793 | 39 | Male | 33.4 | Non-diabetic |
| 15 | SAMN33103085 | 49 | Female | 27.5 | Non-diabetic |
| 16 | SAMN34033793 | 39 | Male | 33.4 | Non-diabetic |
| 17 | UNOSIDAKE4447 | 27 | Male | 29.79 | Type 1 Diabetic (10 years) |
| 18 | SAMN37680485 | 63 | male | 33.9 | Type 2 Diabetic (>10 years) |
| 19 | SAMN36020139 | 46 | Female | 27.5 | Type 2 Diabetic (0-5 years) |
| 20 | SAMN33293779 | 49 | Male | 68.5 | Type 2 Diabetic (Duration unknown) |
| 21 | SAMN25980818 | 55 | Male | 31 | Type 2 diabetic (0-5 years) |
| 22 | SAMN39639181 | 58 | Male | 38.3 | Type 2 diabetic (6-10 years) |
| 23 | SAMN32273466 | 45 | Female | 33.5 | Type 2 diabetic (Duration unknown) |
| 24 | SAMN40122446 | 63 | Male | 36 | Type 2 diabetic (>10 years) |
| 25 | SAMN39243818 | 68 | Male | 25.2 | Type 2 diabetic (>10 years) |
| 26 | SAMN40593976 | 43 | Male | 29.8 | Type 2 diabetic |

**Supplementary Table S2. Antibody information. Related to Figures 2J, 3H, 3N, 4A, 4E and 5D.**

| Target | Primary Antibody (Dilution) | Secondary Antibody (1:1000 dilution) |
|---|---|---|
| Nidogen-2 | Rat Monoclonal Antibody (1:50) | Anti-Rat Alexa fluor 633 |
| GLUT4 | Polyclonal Rabbit GLUT4 Antibody (1:100) | Anti-Rabbit Alexa fluor 488 |
| α-sarcoglycan | Monoclonal mouse a-sarcoglycan Antibody (F-7) (1:100) | Anti-mouse Alexa fluor 594 |
| UCP1 | monoclonal rabbit UCP1 antibody (1:50) | Anti-Rabbit Alexa fluor 488 |
| Insulin | Monoclonal Mouse Insulin Antibody (1:100) | Anti-mouse Alexa fluor 568 |
| Somatostatin | Monoclonal Rat Somatostatin Antibody (1:50) | Anti-Rat Alexa fluor 633 |
| Glucagon | Rabbit GluN F7 (1:100) | Anti-Rabbit Alexa fluor 488 |

**Supplementary Table S3. Primers for qPCR amplification. Related to Figures 3G, 3I, 3M, 3O, 4D, and S2L.**

| Target | Gene Name | Forward primer | Reverse Primer |
|---|---|---|---|
| *Ucp1* | Uncoupling protein 1 | GATGGTGAACCCGACAACTT | AGCACACAAACATGATGACGT |
| *18S* | Small subunit 18 rRNA | GTAACCCGTTGAACCCCATT | CCATCCAATCGGTAGTAGCG |
| *Cidea* | Cell Death Inducing DFFA-Like Effector A | TGCTCTTCTGTATCGCCCAGT | GCCGTGTTAAGGAATCTGCTG |
| *Zic1* | Zic Family Member 1 | AACCTCAAGATCCACAAAAGGA | CCTCGAACTCGCACTTGAA |
| *Prdm16* | PR/SET Domain 16 | CAGCACGGTGAAGCCATTC | GCGTGCATCCGCTTGTG |
| *Rbp4* | Retinol binding protein 4 | TGTAGCCTCCTTTCTCCAGCG | ACAGGTGCCATCCAGATTCTG |
| *Cxcl14* | Chemokine(C-X-C motif) ligand 14 | TACCCACACTGCGAGGAGAAG | CGCTTCTCGTTCCAGGCATTG |
| *Trarg1* | Trafficking regulator of GLUT4-1 | TGGCACGGCTACTCAGCATCA | TGCCTCTGCTAGAAACAGCTCC |
| *Glut4* | Glucose transporter type 4 | GGTGTGGTCAATACGGTCTTCAC | AGCAGAGCCACGGTCATCAAGA |

### Supplementary Table S4. RNAseq Top 15 DEGs from DESeq2 of CB-100 treated versus sham-treated NOD Mouse's ingWAT. Related to Figure 3D.

| Gene Symbol | BaseMean | log2 FoldChange | lfcSE | Stat | pvalue | padj | Note |
|---|---|---|---|---|---|---|---|
| ***Prr32*** | **98.984989** | **9.89392454** | **1.56387026** | **6.32656353** | **2.5068E-10** | **6.4319E-09** | **upregulated** |
| ***Sprr2a3*** | **84.7456472** | **9.67043129** | **1.87952704** | **5.14514082** | **2.6732E-07** | **3.1905E-06** | **upregulated** |
| ***Sprr2f*** | **71.8354655** | **9.43189703** | **1.87675419** | **5.02564324** | **5.0175E-07** | **5.5401E-06** | **upregulated** |
| ***Sprr1a*** | **237.263237** | **7.96854276** | **1.12842095** | **7.06167566** | **1.6451E-12** | **7.0183E-11** | **upregulated** |
| ***Lep*** | **6639.40122** | **7.78243891** | **0.80491122** | **9.66869225** | **4.0958E-22** | **8.8054E-20** | **upregulated** |
| *Slc2a5* | 353.175461 | 6.60632428 | 1.35415758 | 4.87854911 | 1.0687E-06 | 1.0717E-05 | upregulated |
| *Fpr1* | 323.351452 | 6.56658049 | 1.42605491 | 4.60471784 | 4.1303E-06 | 3.4666E-05 | upregulated |
| *Alb* | 450.504986 | 6.43586345 | 0.54351805 | 11.8411217 | 2.3923E-32 | 2.3144E-29 | upregulated |
| *Sncg* | 15023.3037 | 6.23226359 | 0.34513761 | 18.0573299 | 6.9084E-73 | 1.1362E-68 | upregulated |
| *Pnpla3* | 5753.84419 | 6.20018137 | 0.73360627 | 8.45164721 | 2.8722E-17 | 2.9803E-15 | upregulated |
| *Areg* | 1544.70625 | 6.18068349 | 1.2612098 | 4.90059899 | 9.5545E-07 | 9.7209E-06 | upregulated |
| ***Scd1*** | **672756.834** | **6.14968158** | **0.68303094** | **9.00351835** | **2.186E-19** | **3.1957E-17** | **upregulated** |
| *Upk3a* | 107.441587 | 6.04186451 | 1.76080707 | 3.4313041 | 0.00060069 | 0.00262152 | upregulated |
| *Mup10* | 134.373557 | 6.02375742 | 1.1716738 | 5.1411557 | 2.7305E-07 | 3.2518E-06 | upregulated |
| *Mup22* | 133.025014 | 6.00902777 | 1.16948656 | 5.13817599 | 2.7742E-07 | 3.2966E-06 | upregulated |
| *Hamp2* | 65.2044088 | -5.37225207 | 0.74256854 | -7.23468845 | 4.666E-13 | 2.1832E-11 | downregulated |
| *Ighv1-49* | 66.5498465 | -4.50562545 | 0.86899177 | -5.18488852 | 2.1614E-07 | 2.6519E-06 | downregulated |
| *Ighe* | 1259.95447 | -3.86656127 | 0.98426646 | -3.9283684 | 8.5524E-05 | 0.00048245 | downregulated |
| *Igkv11-125* | 65.3703606 | -2.95852355 | 0.7937741 | -3.72716059 | 0.00019365 | 0.00097307 | downregulated |
| *Apoc3* | 59.3517823 | -2.90471797 | 0.68151176 | -4.26216855 | 2.0245E-05 | 0.00013736 | downregulated |
| *A530030E21Rik* | 145.876251 | -2.8646756 | 0.23770678 | -12.0512993 | 1.9092E-33 | 2.0257E-30 | downregulated |
| *Gm18748* | 63.5887462 | -2.72271781 | 0.49317453 | -5.52079975 | 3.3746E-08 | 5.0941E-07 | downregulated |
| *Gm47782* | 203.285799 | -2.71880525 | 0.44753582 | -6.07505621 | 1.2394E-09 | 2.7125E-08 | downregulated |
| *Gm31804* | 270.517933 | -2.70462141 | 0.42806294 | -6.3182797 | 2.6449E-10 | 6.7389E-09 | downregulated |
| *Abca13* | 55.4327726 | -2.6672388 | 0.48798094 | -5.46586673 | 4.6065E-08 | 6.7283E-07 | downregulated |
| *Ptgds* | 62.7145289 | -2.65517546 | 0.49570735 | -5.35633664 | 8.4926E-08 | 1.163E-06 | downregulated |
| *Gm12490* | 63.6375533 | -2.62447864 | 0.42457837 | -6.18137625 | 6.3545E-10 | 1.4814E-08 | downregulated |
| *Gm47780* | 241.250346 | -2.62399536 | 0.46867817 | -5.59871464 | 2.1595E-08 | 3.4431E-07 | downregulated |
| *Olfr109* | 217.333967 | -2.49909268 | 0.37270234 | -6.70533142 | 2.0095E-11 | 6.7724E-10 | downregulated |
| *Mir142b* | 76.7799563 | -2.48147157 | 0.45053798 | -5.50779663 | 3.6335E-08 | 5.4475E-07 | downregulated |

Noteworthy genes are highlighted in red.

BaseMean = the average of the normalized counts taken over all samples / lfcSE = standard error of the log2FoldChange estimate/stat = Wald statistic / pvalue = Wald test p-value / padj = Benjamini-Hochberg adjusted p-value[1].

**Supplementary Table S5. RNAseq Top 15 DEGs from DESeq2 of CB-100 treated versus sham-treated NOD Mouse's BAT. Related to Figure 3J.**

| Gene Symbol | BaseMean | log2 FoldChange | lfcSE | stat | pvalue | padj | Note |
|---|---|---|---|---|---|---|---|
| *Fosb* | 377.940493 | 5.423568265 | 1.630662935 | 3.325989785 | 0.000881052 | 0.014228774 | upregulated |
| ***Scd1*** | **43378.4807** | **4.404576517** | **0.502375191** | **8.767504048** | **1.8267E-18** | **4.4462E-15** | **upregulated** |
| *Egr3* | 193.339786 | 4.170345034 | 1.138303094 | 3.663650792 | 0.000248646 | 0.005892108 | upregulated |
| ***Gm44502*** | **147.105008** | **3.430936505** | **0.600836449** | **5.710266926** | **1.12799E-08** | **2.66927E-06** | **upregulated** |
| ***Dio2*** | **1536.95743** | **3.073423038** | **1.144592535** | **2.685167816** | **0.00724934** | **0.060724803** | **upregulated** |
| *Gm38357* | 480.556497 | 3.009943113 | 0.745179096 | 4.039221079 | 5.3629E-05 | 0.001887874 | upregulated |
| *Fos* | 424.388827 | 2.951669149 | 1.39495136 | 2.115965642 | 0.03434773 | 0.16517545 | upregulated |
| *Slco4a1* | 133.237886 | 2.825701883 | 0.963917187 | 2.931477849 | 0.003373534 | 0.036218196 | upregulated |
| *Fndc5* | 65.938467 | 2.644053492 | 0.656213661 | 4.029257007 | 5.59534E-05 | 0.001957565 | upregulated |
| *Akap5* | 142.150916 | 2.628803173 | 0.313919397 | 8.374134244 | 5.56313E-17 | 1.05316E-13 | upregulated |
| *Gk* | 1773.065 | 2.573316009 | 0.462775949 | 5.560608789 | 2.68835E-08 | 5.45287E-06 | upregulated |
| *A830018L16Rik* | 183.331672 | 2.515879267 | 0.52175858 | 4.82192218 | 1.42181E-06 | 0.000114015 | upregulated |
| *S100b* | 336.242184 | 2.490030416 | 0.423136754 | 5.884694227 | 3.98791E-09 | 1.23538E-06 | upregulated |
| *Gm43605* | 122.466743 | 2.467060482 | 0.561606957 | 4.392859542 | 1.11869E-05 | 0.000552473 | upregulated |
| *Gm42613* | 51.4196396 | 2.441051139 | 0.56734788 | 4.302565015 | 1.68832E-05 | 0.000775286 | upregulated |
| *Gm45061* | 548.29461 | -3.843311266 | 0.486025033 | -7.90764057 | 2.62313E-15 | 4.06299E-12 | downregulated |
| *Otof* | 145.001663 | -3.829071067 | 1.257713583 | -3.044469837 | 0.002330908 | 0.027768239 | downregulated |
| *Cyp2b10* | 809.7713 | -3.778733581 | 0.762610567 | -4.954997669 | 7.23312E-07 | 6.52052E-05 | downregulated |
| *Chrnb4* | 52.0726661 | -3.728396563 | 1.523326905 | -2.447535424 | 0.014383699 | 0.094914584 | downregulated |
| *Itih4* | 137.694373 | -3.51309137 | 0.902616415 | -3.892119967 | 9.93721E-05 | 0.002982145 | downregulated |
| *Zim1* | 682.570621 | -3.441861698 | 0.478980903 | -7.185801513 | 6.68141E-13 | 7.11486E-10 | downregulated |
| *Gm45060* | 132.597611 | -3.383178152 | 0.358909528 | -9.426270106 | 4.24931E-21 | 2.41333E-17 | downregulated |
| *Gm16143* | 60.2954912 | -3.222388937 | 0.462843773 | -6.962152515 | 3.35112E-12 | 2.48245E-09 | downregulated |
| *BB365896* | 1183.13087 | -3.047122329 | 0.774020225 | -3.936747687 | 8.25934E-05 | 0.002586812 | downregulated |
| *Mycl* | 297.128494 | -2.999406415 | 0.550259653 | -5.450892858 | 5.01176E-08 | 8.63586E-06 | downregulated |
| *D630039A03Rik* | 68.4987102 | -2.973999314 | 1.189594744 | -2.500010469 | 0.012418964 | 0.086271408 | downregulated |
| *Cpne9* | 182.564779 | -2.824471637 | 0.495642026 | -5.698612077 | 1.20787E-08 | 2.78103E-06 | downregulated |
| *Tmem132b* | 125.969361 | -2.797054301 | 0.90575213 | -3.088101266 | 0.002014398 | 0.025219015 | downregulated |
| *Hmgcs2* | 154.102529 | -2.716866552 | 0.602274369 | -4.51101141 | 6.45192E-06 | 0.000382662 | downregulated |
| *Aldoc* | 90.026831 | -2.696693801 | 0.290174448 | -9.293353778 | 1.49502E-20 | 5.09442E-17 | downregulated |

Noteworthy genes are highlighted in red.

BaseMean = the average of the normalized counts taken over all samples / lfcSE = standard error of the log2FoldChange estimate/stat = Wald statistic / pvalue = Wald test p-value / padj = Benjamini-Hochberg adjusted p-value[1].
